# ORMDL3 restrains type-I interferon signaling and anti-tumor immunity by promoting RIG-I degradation

**DOI:** 10.1101/2024.08.13.607743

**Authors:** Qi Zeng, Chen Yao, Shimeng Zhang, Yizhi Mao, Jing Wang, Ziyang Wang, Chunjie Sheng, Shuai Chen

**Affiliations:** State Key Laboratory of Oncology in South China, Guangdong Provincial Clinical Research Center for Cancer, Sun Yat-sen University Cancer Center, Guangzhou 510060, P. R. China; Center for Translational Medicine, The First Affiliated Hospital, Sun Yat-sen University, Guangzhou, Guangdong, 510080, P.R.China

**Keywords:** ORMDL3, IFN, RIG-I, antiviral innate immunity, anti-tumor immunity

## Abstract

Mounting evidence has demonstrated the genetic association of ORMDL3 (ORMDL Sphingolipid Biosynthesis Regulator 3) gene polymorphisms with bronchial asthma and a diverse set of inflammatory disorders. However, its role in type I interferon (IFN) signaling remains poorly defined. Herein, we report that ORMDL3 is a negative modulator of the type I IFN signaling by interacting with MAVS (Mitochondrial Antiviral Signaling protein) and subsequently promoting the proteasome-mediated degradation of RIG-I (Retinoic Acid-Inducible Gene I). Immunoprecipitation coupled with mass spectrometry (IP-MS) assays uncovered that ORMDL3 binds to USP10 (Ubiquitin-Specific Protease 10), which forms a complex with and stabilizes RIG-I through decreasing its K48-linked ubiquitination. ORMDL3 thus disrupts the interaction between USP10 and RIG-I, thereby promoting RIG-I degradation. Additionally, subcutaneous syngeneic tumor models in C57BL/6 mice revealed that inhibition of ORMDL3 enhances anti-tumor efficacy by augmenting the proportion of cytotoxic CD8 positive T cells and IFN production in the tumor microenvironment (TME). Collectively, our findings reveal the pivotal roles of ORMDL3 in maintaining antiviral innate immune responses and anti-tumor immunity.

## Introduction

Type I interferons (IFN-Is) play a key role in the innate immune response to viral infections. Under viral stimulations, cells produce and release interferons, which induce the transcription of interferon-stimulated genes ^1,2^. Besides their critical role in antiviral immune responses, growing evidence suggests that IFN-Is produced by malignant tumor cells or infiltrating immune cells also influence the effectiveness of cancer immunotherapy^3–5^. Many traditional chemotherapeutic drugs, targeted anti-tumor drugs, immunoadjuvants, and oncolytic viruses require intact type I IFN signaling to exert their full effects ^6,7^. Furthermore, studies have shown that high intratumoral expression levels of IFN-Is or IFN-stimulated genes are associated with positive disease outcomes in cancer patients^8^.

In response to viral RNA molecules, RLRs (RIG-I-like receptors) exposed its CARD domain and then cooperate with mitochondrial antiviral signaling protein (MAVS) and TANK-binding kinase1(TBK1) to promote the production of IFN-Is^9^. TBK1 phosphorylates and activates IRF3 and IRF7, which then induce the expression of IFN-Is and various interferon-stimulated genes^10,11^. Thus, targeting RLRs can provoke anti-infection activities, moreover, RLRs also play an important role in anti-tumor immunity, for example, targeting RLRs can sensitize “immune-cold” tumors to become “immune hot” tumors^12^. DNA methyltransferase inhibitors upregulate endogenous retroviruses in tumor cells to induce the activation of the RLR-mediated RNA recognition pathway that potentiates immune checkpoint therapy^13,14^. SB 9200 (also known as inarigivir soproxil or GS 9992) is an orally available prodrug of a dinucleotide agonist of RIG-I and nucleotide binding oligomerization domain-containing protein 2 (NOD2)^15^ and is currently in clinical trials to treat chronically infected HCV patients^16^. Another RIG-I agonist, MK-4621, appears to be safe and well-tolerable for patients with advanced or recurrent tumors, with no dose-limiting toxicities^17^.

The post-transcriptional modifications (PTMs) of RIG-I are vital for its activation and stability. Several E3-ligases have been reported to catalyze K63- or K48-linked polyubiquitination of RIG-I, regulating the RLR pathway. K63-linked ubiquitination,mediated by TRIM25^18^ and Mex-3 RNA binding family member C (MEX3C) in the CARD region of RIG-I^19,20^, or by RNF135 in the C-terminus of RIG-I^21,22^, facilitates RLR signal transduction. In addition to K63 ubiquitination that usually associates with signal transduction pathways, classical degradative K48-linked polyubiquitylation also regulates RIG-I’s stability. Several E3 ligases such as Cbl^23^, ring finger protein 122 (RNF122)^24^, RNF125^25^ and TRIM40^26^, catalyze this process. Conversely, several deubiquitylating enzymes also regulate RIG-I expression. For example, ubiquitin-specific peptidase 3 (USP3), USP21 and CYLD lysine 63 deubiquitinase (CYLD) modulate RIG-I signaling by removing K63-polyubiquitin chains^27,28^. Deubiquitinase USP4 and USP15 can increase the stability of RIG-I and TRIM25 by decreasing K48 ubiquitination of them^29,30^. Discovering new proteins regulating the activity or stability of RIG-I will provide new insights and targets for antiviral and anti-tumor therapies.

ORMDL3 is a member of the three-gene ORDML family (ORMDL1, ORMDL2, and ORMDL3), and it is a 153-aa transmembrane protein primarily located in the endoplasmic reticulum (ER)^37^. Genetic variants in *ORMDL3* are associated with sphingolipid synthesis and altered metabolism, which contribute to asthma^31^. Recent evidence has elucidated that ORMDL3 regulates eosinophil trafficking, recruitment, and degranulation, which may induce the formation of allergic asthma and potentially other eosinophilic disorders^32^. Additionally, ORMDL3 polymorphisms also contribute to a diverse set of inflammatory disorders that include bronchial asthma, inflammatory bowel disease^33^, ankylosing spondylitis^34^, atherosclerosis^35^, SLE^36^ and cholangitis^37,38^. However, the role of ORMDL3 in innate immunity remains unknown.

In this study, we illuminate ORMDL3 as a pivotal negative regulator of the type I interferon (IFN) signaling pathway. ORMDL3 forms a complex with MAVS and subsequently directs RIG-I toward degradation. ORMDL3 amplifies the K48-linked ubiquitination of RIG-I by disrupting the interaction between RIG-I and USP10. Animal experiments showed that inhibiting ORMDL3 enhances anti-tumor activity, demonstrated by an augmented proportion of activated CD8^+^ T cells and increased interferon production within the tumor microenvironment (TME). Collectively, our results unveil the critical role of ORMDL3 in maintaining the homeostasis of antiviral innate immune responses and suggest ORMDL3 as a candidate target for cancer immunotherapy.

## Results

### ORMDL3 negatively regulates RLR induced type I IFN signaling pathway

In order to investigate the potential role of ORMDL3 in the antiviral response, HEK293T cell overexpression ORMDL3 was stimulated with poly(I:C) or VSV infection. The result showed that ORMDL3 significantly inhibited poly(I:C) and VSV stimulated transcription of *IFNB1* **(Figure 1A)**. Western blots demonstrated a marked reduction in the phosphorylation level of IRF3 when ORMDL3 was overexpressed **(Figure 1B**). Ectopic expressed ORMDL3 in A549 cells attenuated *IFNB1* expression induced by poly(I:C) but not poly(dG:dC) **(Figure 1C)**. This phenomenon was also observed in parallel experiments using primary mouse BMDM cells **(Figure 1D**). These findings highlight ORMDL3 as a repressor of RNA-induced type I IFN expression. To unravel the molecular mechanism underlying the suppression of type I IFN signaling by ORMDL3, luciferase reporter assays were performed. ORMDL3 was found to decrease the luciferase reporter activity induced by RIG-I, while showing no effect on cGAS/STING or TRIF **(Figure 1E)**. The ISRE luciferase reporter assay also showed similar result **(Figure 1F)**. We constructed an ORMDL3 stable knockdown cell line of A549 to examine the role of ORMDL3 on viral replication. We found that ORMDL3 knockdown strikingly suppressed the replication of vesicular stomatitis virus (VSV), and overexpression ORMDL3 enhanced the replication of VSV **(Figure 1G)**. We also infected the A549 stable knockdown cell line with herpes simplex virus-1 (HSV-1), and there is no significance between shNC and shORMDL3 cell lines **(Figure 1H)**. These results suggest that ORMDL3 only facilitates RNA virus replication but not DNA virus, which coincides with the finding that ORMDL3 specifically represses RNA but not DNA-induced type I interferon expression (**Figure 1D**).

**Figure 1.**
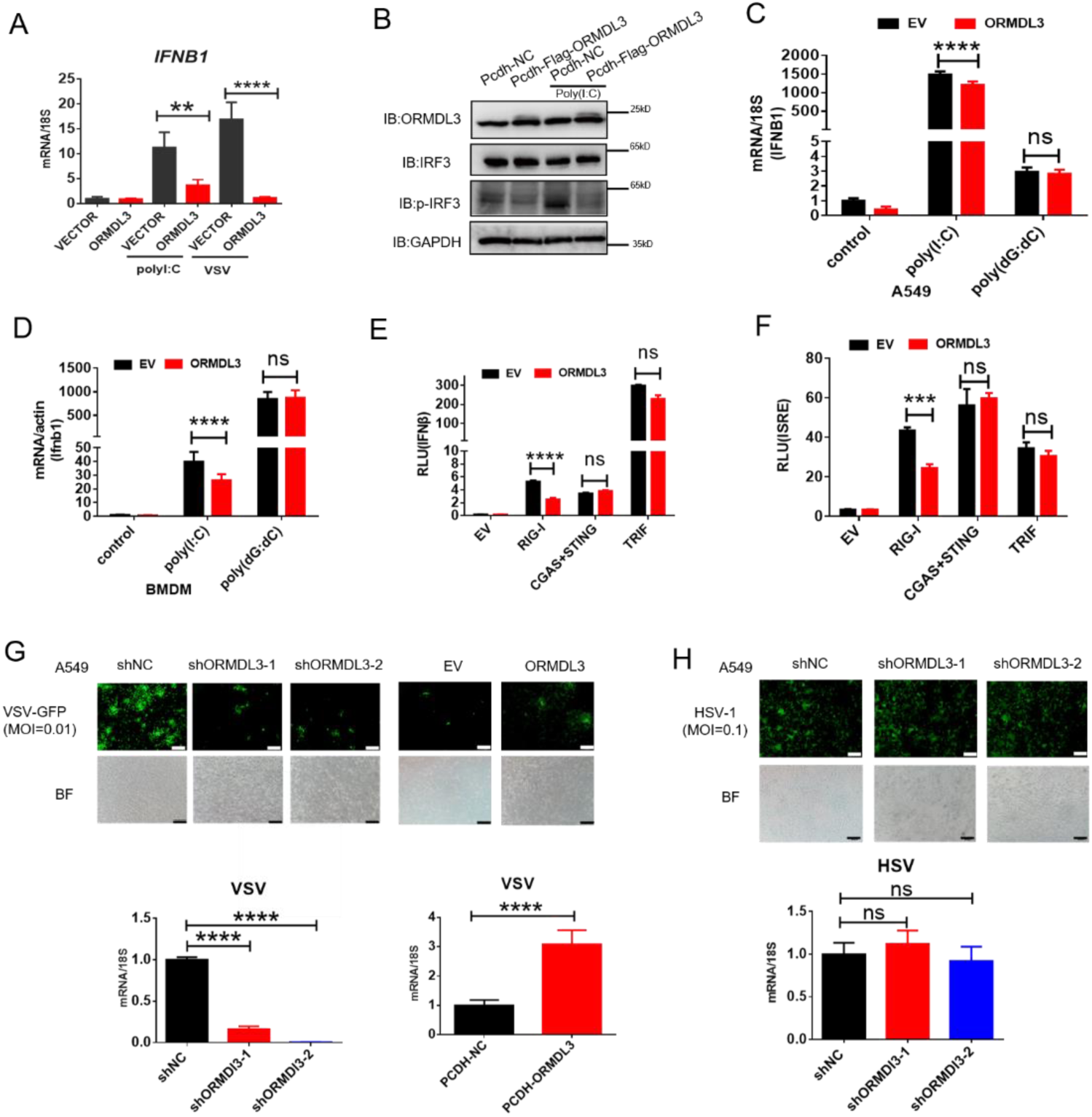
ORMDL3 negatively regulates RLR induced type I IFN signaling pathway. **(A)** HEK293T cells were transfected with an empty vector (EV) or a ORMDL3 plasmid for 12 h and were then infected with VSV (MOI = 0.01) or transfected with poly(I:C). The expression of the mRNAs encoded by the indicated genes was detected using qRT-PCR. **(B)** HEK293T -EV and HEK293T–Flag-ORMDL3 cells were transfected with or without poly(I:C), and immunoblot analyses of phosphorylated IRF3 (p-IRF3), total IRF3, GAPDH and ORMDL3 levels were performed. **(C)** Results of the qRT-PCR assays showing mRNA levels of *IFNB1* in A549 cells transfected with control or ORMDL3 followed by transfected with poly(I:C) or poly(dG:dC). **(D)** Results of the qRT-PCR assays showing mRNA levels of *Ifnb1* in mice primary BMDM cells infected with control or psc-AAV-ORMDL3 virus followed by transfected with poly(I:C) or poly(dG:dC). **(E)** Results of the luciferase assay showing IFN-β-Luc activity in HEK293T cells transfected with EV or ORMDL3 plasmids together with individual EV, RIG-I, cGAS plus STING, and TRIF plasmids for 24 h. **(F)** Results of the luciferase assay showing ISRE-Luc activity in HEK293T cells transfected with EV or ORMDL3 plasmids together with individual EV, RIG-I, cGAS plus STING, and TRIF plasmids for 24 h. **(G)** Wild-type and ORMDL3 stable knockdown and overexpression A549 cells were infected with VSV-GFP (MOI = 0.01) for 12 hours, and viral infectivity was detected using fluorescence microscopy, viral amount was detected using RT-PCR, scale bars, 200μm. **(H)** Wild-type and ORMDL3 stable knockdown A549 cells were infected with HSV-1 (MOI = 0.1) for 24 h, and viral infectivity was detected using fluorescence microscopy, viral amount was detected using RT-PCR, scale bars, 200μm. *p < 0.05, **p < 0.01, ****p < 0.0001. Data are representative of at least two independent experiments.

We next evaluated the influence of VSV, HSV-1 and RIG-I agonist SB9200 on ORMDL3 expression. Given the single nucleotide polymorphism (SNP) site rs7216389 at ORMDL3 locus is associated with the susceptibility of childhood asthma^39^ and virus-induced respiratory wheezing illnesses ^40^, we took the genotype of this SNP into account. Upon these stimulations, HSV-1 does not obviously alter the abundance of ORMDL3 **(Figure S1A)**. For VSV and SB920, the expression of ORMDL3 is downregulated in some cell lines and is independent of the SNP **(Figure S1B, S1C)**. Taken together, these results suggest that ORMDL3 is a negative regulator of RLR RNA sensing pathway, and its expression is reciprocally repressed by this pathway.

### ORMDL3 regulates the protein abundance of RIG-I

Further RT-PCR results indicated that ectopic expression of ORMDL3 inhibited *IFNB1* mRNA expression and transcription of downstream genes *CCL5* and *CXCL10* induced by RIG-I and MAVS but not TBK1 or IRF3-5D **(Figure 2A,2B, S2A)**. Additionally, MDA5-induced IFN upregulation was also inhibited by ORMDL3 **(Figure S2B)**. These results revealed that ORMDL3 negatively regulates RLR pathway. Owing to ORMDL3 was down-regulated in response to VSV stimulation in HEK293T, so we firstly transfected cells with siORMDL3 followed by secondary transfection with RIG-I-N (an active form of RIG-I) in HEK293T, and we found that siORMDL3 significantly increase the expression of *IFNB1*, *CCL5* and *ISG54* **(Figure 2C)** as well as the protein abundance of RIG-I **(Figure 2D)**. As ORMDL3 antibody can recognize all ORMDL family members (ORMDL1, 2 and 3), so we also detected the mRNA level of ORMDL3 to further validate knock-down efficiency **(Figure S2C)**. Subsequent experiments involving various signaling proteins such as RIG-I (WT/N), MDA5, TBK1, and IRF3 indicated a negative correlation between ORMDL3 levels and RIG-I/RIG-I-N protein expression, with maximal degradation observed in RIG-I-N **(Figure 2E)**. To investigate that whether ORMDL3 is unique in promoting RIG-I degradation, we compared ORMDL1, 2 and 3, when we co-expresssed RIG-I-N with them, we found that only ORMDL3 can facilitate the degradation of RIG-I-N **(Figure S2D)**. In addition, we also tested whether overexpressing mice Ormdl3 will also lead to mice Rig-i degradation. Interestingly we discovered that mice Rig-I and human RIG-I both can be degraded upon ORMDL3 overexpression, implying that ORMDL3’s function is conservative in human and mice (**Figure S2E,F**). In addition, ORMDL3 overexpression also eliminated endogenous RIG-I protein abundance **(Figure S2G)**.

**Figure 2.**
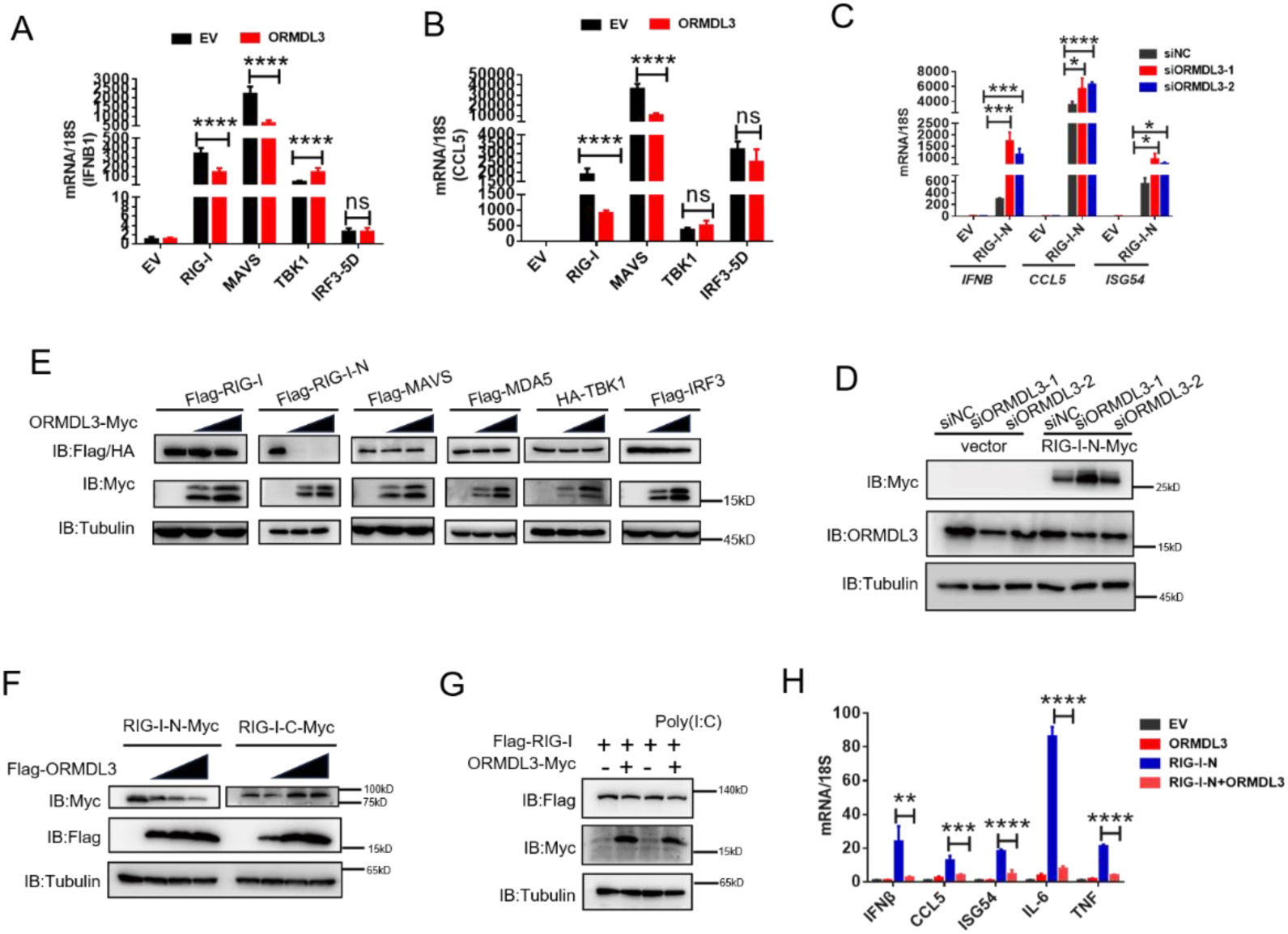
ORMDL3 regulates the protein abundance of RIG-I. **(A,B)** RT-PCR analyses of the expression of the indicated mRNAs in HEK293T cells transfected with EV or ORMDL3 plasmids combined with co-transfection of individual plasmids encoding EV, RIG-I, MAVS, TBK1,IRF3-5D. **(C)** Results of the qRT-PCR assays showing mRNA levels of IFNB1 CCL5 and ISG54 in HEK293T cells transfected with control or ORMDL3-specific siRNAs followed by secondary transfection with EV or RIG-I-N plasmids. **(D)** Results of the WB assays show protein levels in HEK293T cells transfected with control or ORMDL3-specific siRNAs followed by secondary transfection with EV or RIG-I-N plasmids. **(E)** Immunoblot analysis of extracts of 293T cells transfected with expression vector for Flag tagged RIG-I, RIG-I-N, MAVS, MDA5, IRF3 and HA-TBK1, and increasing doses of expression vector for ORMDL3-Myc. **(F)** Immunoblot analysis of extracts of 293T cells transfected with expression vector for RIG-I-N-Myc and RIG-I-C-Myc and increasing doses of expression vector for Flag-ORMDL3. **(G)** HEK293T cells were transfected with Flag-RIG-I and ORMDL3-Myc plasmids, as indicated, with or without poly(I:C) co-transfection. Cell lysates were immunoblotted with a-Flag and a-Myc antibodies. **(H)** Results of the qRT-PCR.assays showing IFNB1, CCL5, ISG54, IL-6 and TNF mRNA in HEK293T cells transfected with EV or ORMDL3 plasmids together with individual RIG-I-N plasmids for 24 h. *p < 0.05, **p < 0.01, ****p < 0.0001. Data are representative of at least two independent experiments.

Based on these observations, we focus on the relationship between RIG-I and ORMDL3. We ectopically expressed an increasing amount of Flag-ORMDL3 with the RIG-I-N (harbors two CARD domains) and the RIG-I-C truncated form **(Figure S2H)** and found that ORMDL3 only decreased the protein abundance of RIG-I-N **(Figure 2F).** We co-transfected ORMDL3 and RIG-I with or without poly(I:C) and found that RIG-I was degraded upon the stimulation of poly(I:C), suggesting that ORMDL3 degrades RIG-I only when it was activated and the CARD domain was exposed **(Figure 2G)**. Further examination showed that co-expression of ORMDL3 suppressed RIG-I-N induced expression of *IFNB1* and *CCL5*, as well as pro-inflammatory cytokines *IL-6* and *TNF* **(Figure 2H)**, suggesting its role on both NF-κB and type I interferon pathways. Given the regulation of ORMDL3 on NF-κB has been reported, we focused on its role in the type I interferon pathway.

### ORMDL3 promotes the proteasome degradation of RIG-I

Two major protein degradation pathways including the ubiquitin proteasome pathway and lysosoml proteolysis system. We next identified which degradation system dominantly mediates the degradation of RIG-I by ORMDL3. We co-expressed ORMDL3 and RIG-I and treated cells with proteasome inhibitor MG132, and lysosome inhibitor CQ. We found the degradation of RIG-I mediated by ORMDL3 could be blocked by proteasome inhibitor MG132 but not lysosome inhibitor CQ **(Figure 3A)**. To rule out the possibility of transcriptional down-regulation of RIG-I, RT-PCR analysis was performed, confirming that the decrease in RIG-I protein was a post-transcriptional event **(Figure S3A)**. To investigate the mechanism, we co-transfected RIG-I-N, ORMDL3 and plasmids encoding different forms of ubiquitin. The results indicated an increase in K48-linked ubiquitin chains on RIG-I-N, implicating ORMDL3 in promoting proteasomal degradation of RIG-I **(Figure 3B)**. To pinpoint the lysine residues crucial for RIG-I ubiquitination, we engineered a mutant version of RIG-I-N in which all lysines were mutated to arginines, denoted as RIG-I-N-KR. Intriguingly, the degradation-promoting effect of ORMDL3 on RIG-I-N-KR was nullified **(Figure 3C)**. Given there are 18 lysine residues in RIG-I-N, we generated two mutants, mutant1 and mutant2, each mutating the last nine lysines and the first nine lysines **(Figure S3B)**. Remarkably, the results showed that mutant1 was resistant to degradation by ORMDL3 **(Figure 3D)**, suggesting that the last nine lysines on RIG-I-N mediated its degradation induced by ORMDL3.

**Figure 3.**
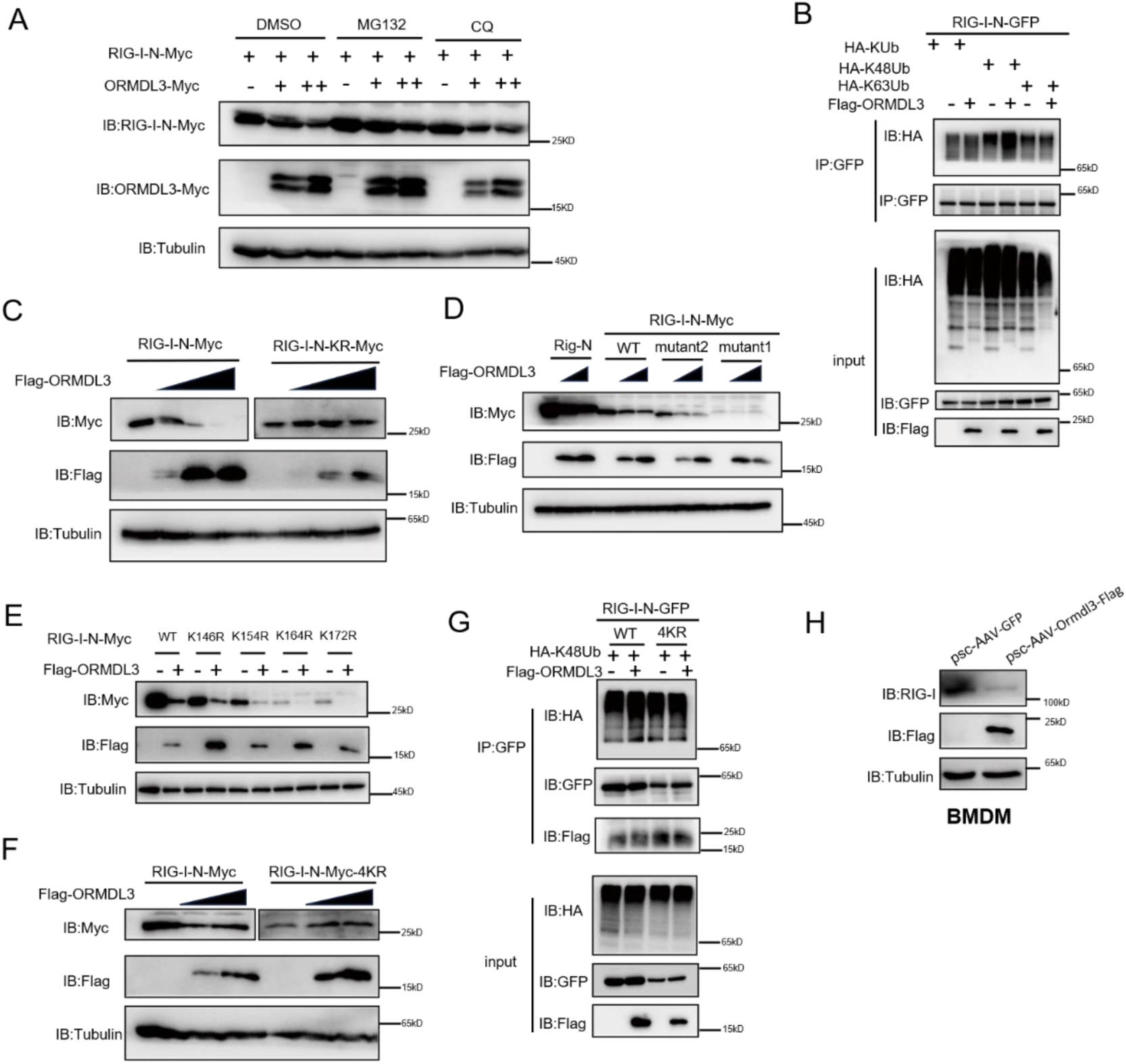
ORMDL3 promotes the proteasome degradation of RIG-I. **(A)** HEK293T cells were transfected with plasmids encoding RIG-I-N-Myc together with increasing amounts of Flag-ORMDL3 plasmid treated with MG132 (10 μM), chloroquine (CQ) (50 μM) for 6h and the cell lysates were analyzed by immunoblot. **(B)** HEK293T cells were transfected with the indicated plasmids, and cell lysates were immunoprecipitated with an GFP antibody (a-GFP) followed by immunoblots using GFP (a-GFP) and a-HA antibodies. **(C)** 293T cells were transfected with RIG-I-N-Myc (WT or KR) and increasing doses of expression vector for Flag-ORMDL3. The expression levels of RIG-I-N-Myc were analyzed by immunoblot. **(D)** 293T cells were transfected with Rig—N,RIG-I-N-Myc (WT, KR,mutant1,mutant2), and increasing doses of expression vector for Flag-ORMDL3. The expression levels of RIG-I-N-Myc were analyzed by immunoblot. **(E)** 293T cells were transfected with RIG-I-N-Myc (WT, K146R, K154R, K164R, K172R) with or without Flag-ORMDL3. The expression levels of RIG-I-N-Myc and its mutant forms were analyzed by immunoblot. **(F)** HEK293T cells were transfected with RIG-I-N-Myc (WT or 4KR) and increasing doses of expression vector for Flag-ORMDL3. The expression levels of RIG-I-N-Myc (WT or 4KR) were analyzed by immunoblot. **(G)** HEK293T cells were transfected with RIG-I-N-GFP and HA-KUb,HA-K48Ub and HA-K63Ub with EV or Flag-ORMDL3, and cell lysates were immunoprecipitated with GFP antibody (a-GFP) followed by immunoblots using GFP (a-GFP) and a-HA, a-Flag antibodies. **(H)** BMDM cells were infected with psc-AAV-GFP and psc-AAV-ORMDL3-Flag virus, followed by immunoblot analysis of RIG-I, Flag, and Tubulin.

To delve deeper into the intricate mechanism of ORMDL3-induced degradation of RIG-I, we initially introduced single-point mutations, specifically K146R, K154R, K164R, and K172R ^9^, which have been reported important for the function and stability of RIG-I. Co-transfection with ORMDL3 revealed that these individual mutations did not impede the degradation process, hinting at the potential cooperation of lysine residues **(Figure 3E)**. We then mutated all four lysine residues and assessed ORMDL3-induced RIG-I-N degradation. Strikingly, the RIG-I-N-4KR mutant, in which K146, K154, K164, and K172 were simultaneously mutated to arginines, displayed resistance to degradation by ORMDL3 **(Figure 3F)**. At the meantime, the 4KR mutant failed to exhibit the upregulation of K48-linked ubiquitination induced by ORMDL3 overexpression, reinforcing the pivotal role played by K146, K154, K164 and K172 in mediating RIG-I ubiquitination and subsequent degradation **(Figure 3G)**. In addition, we found that in mice primary BMDM when ORMDL3 was overexpressed, endogenous RIG-I was downregulated **(Figure 3H)**.

### ORMDL3 interacts with the signaling adaptor MAVS

Next, we sought to determine the binding partner of ORMDL3 in the type I interferon pathway. Co-immunoprecipitation(co-IP) and immunoblot analysis showed that only Flag-tagged MAVS interacted with ORMDL3-GFP **(Figure 4A)**. To delineate the requisite domains of MAVS facilitating this interaction, various MAVS truncations were co-transfected with ORMDL3. Notably, deletion of the transmembrane domain (TM) of MAVS abrogated the interaction, while deletion of the caspase activation and recruitment domain (CARD) had no discernible impact **(Figure 4B, S4A)**. It has been reported that ORMDL3 contains four TM segments (TM1−TM4) ^41^. To map the essential domains of ORMDL3 that mediate its association with MAVS, we generated different truncations of ORMDL3 based on the structure, which includes four truncations 1-42, 43-82, 83-118, 119-153 **(Figure S4B)**. Intriguingly, we found that all these four ORMDL3 truncations interact with MAVS **(Figure 4C)** and impede RIG-I-N induced transcription of *IFNB1* and ISGs **(Figure 4D)**. In addition, co-transfection of RIG-I-N-Myc with individual ORMDL3 truncations showed that each domain of ORMDL3 is favorable to RIG-I-N degradation **(Figure 4E)**. Moreover, inspired by Guo et al.’s discovery of a naturally occurring short isoform of ORMDL3^42^, we engineered N- and C-terminal truncations of ORMDL3 to mimic this isoform **(Figure S4B)**, results revealing that both truncations retained the ability to interact with MAVS **(Figure S4C)**. Luciferase assays and RT-PCR affirmed the inhibitory efficacy of each domain **(Figure S4D, E)**. We also tested whether these different domains can promote the degradation of RIG-I-N, results revealed that every domain can amplify the degradation of RIG-I **(Figure S4F)**. To further validate the association between ORMDL3 and MAVS, we did FRET experiment in Hela cells, we overexpressed YFP-MAVS(donor) and CFP-ORMDL3(acceptor), when we bleached YFP-MAVS, and we noticed that the fluorescence of CFP-ORMDL3 enhanced **(Figure 4F)**.

**Figure 4.**
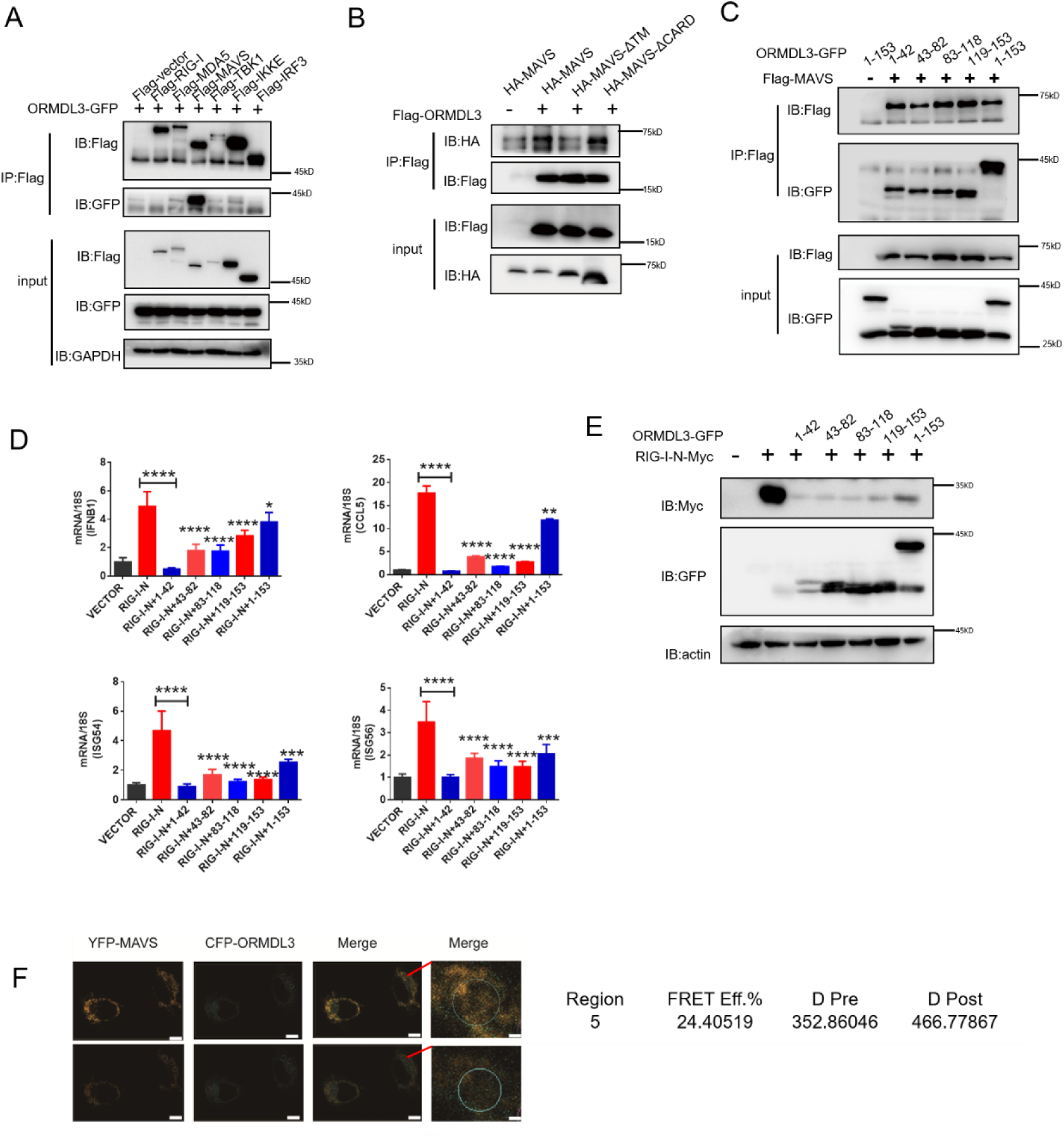
ORMDL3 interacts with signaling adaptor MAVS. **(A)** HEK293T cells were transfected with EV or Flag-RIG-I/MAVS/MDA5/TBK1/IRF3/IKKE with ORMDL3-GFP, and cell lysates were immunoprecipitated with Flag antibody (a-Flag) followed by immunoblots using GFP (a-GFP) and a-Flag antibodies. **(B)** HEK293T cells were transfected with different MAVS truncations with EV or Flag-ORMDL3, and cell lysates were immunoprecipitated with Flag antibody (a-Flag) followed by immunoblots using HA (a-HA) and a-Flag antibodies. **(C)** HEK293T cells were transfected with EV or Flag-MAVS with different ORMDL3 truncations, and cell lysates were immunoprecipitated with Flag antibody (a-Flag) followed by immunoblots using GFP (a-GFP) and a-Flag antibodies. **(D)** HEK293T cells were transfected with EV or Flag-MAVS with different ORMDL3 truncations, followed by RT-PCR analysis of *IFNB1*,*CCL5*,*ISG54*,*ISG56*. **(E)** HEK293T cells were transfected with RIG-I-N-Myc with EV or different ORMDL3 truncations, followed by immunoblots using GFP (a-GFP) and a-Myc antibodies. **(F)** FRET experiment of YFP-MAVS and CFP-ORMDL3 in Hela cells,YFP-MAVS is the donor and CFP-ORMDL3 is the acceptor, FRET efficiency is 24.4019%, scale bars, 10μm. *p < 0.05, **p < 0.01, ****p < 0.0001. Data are representative of at least two independent experiments.

### USP10 deubiquitinates and stablizes RIG-I

To identify the E3 ligase or deubiquitinase involved in ORMDL3-mediated RIG-I ubiquitination, we conducted immunoprecipitation-mass spectrometry (IP-MS) analysis using Flag-ORMDL3 as bait and identified USP10 as a potential candidate **(Figure 5A)**. We validated the IP-MS results and the co-IP experiment revealed that only USP10 can interact with ORMDL3 but not CAND1 or UFL1**(Figure S5)**. Subsequent co-IP validation demonstrated the interaction between ORMDL3 and USP10, USP10 and RIG-I, respectively **(Figure 5B, 5C)**. Interestingly, we found that the RIG-I level is decreased in USP10 stable knockdown 293T cells while overexpression of USP10 promotes the accumulation of RIG-I **(Figure 5D,5E)**. As USP10 is a deubiquitinase, we investigated its impact on RIG-I ubiquitination and observed a decrease in K48 ubiquitination of RIG-I upon USP10 overexpression **(Figure 5F)**. Co-expression RIG-I-N or its 4KR mutant with USP10 showed that USP10 failed to increase the RIG-I-N-4KR level, underscoring the indispensability of these residues **(Figure 5G)**. Upon overexpressing RIG-I-N and ORMDL3 in USP10 knockdown cells, ORMDL3’s ability to degrade RIG-I was markedly compromised, emphasizing the indispensable role of USP10 in this degradation process **(Figure 5H)**. This further validates that ORMDL3 disturbs the USP10’s function on RIG-I.

**Figure 5.**
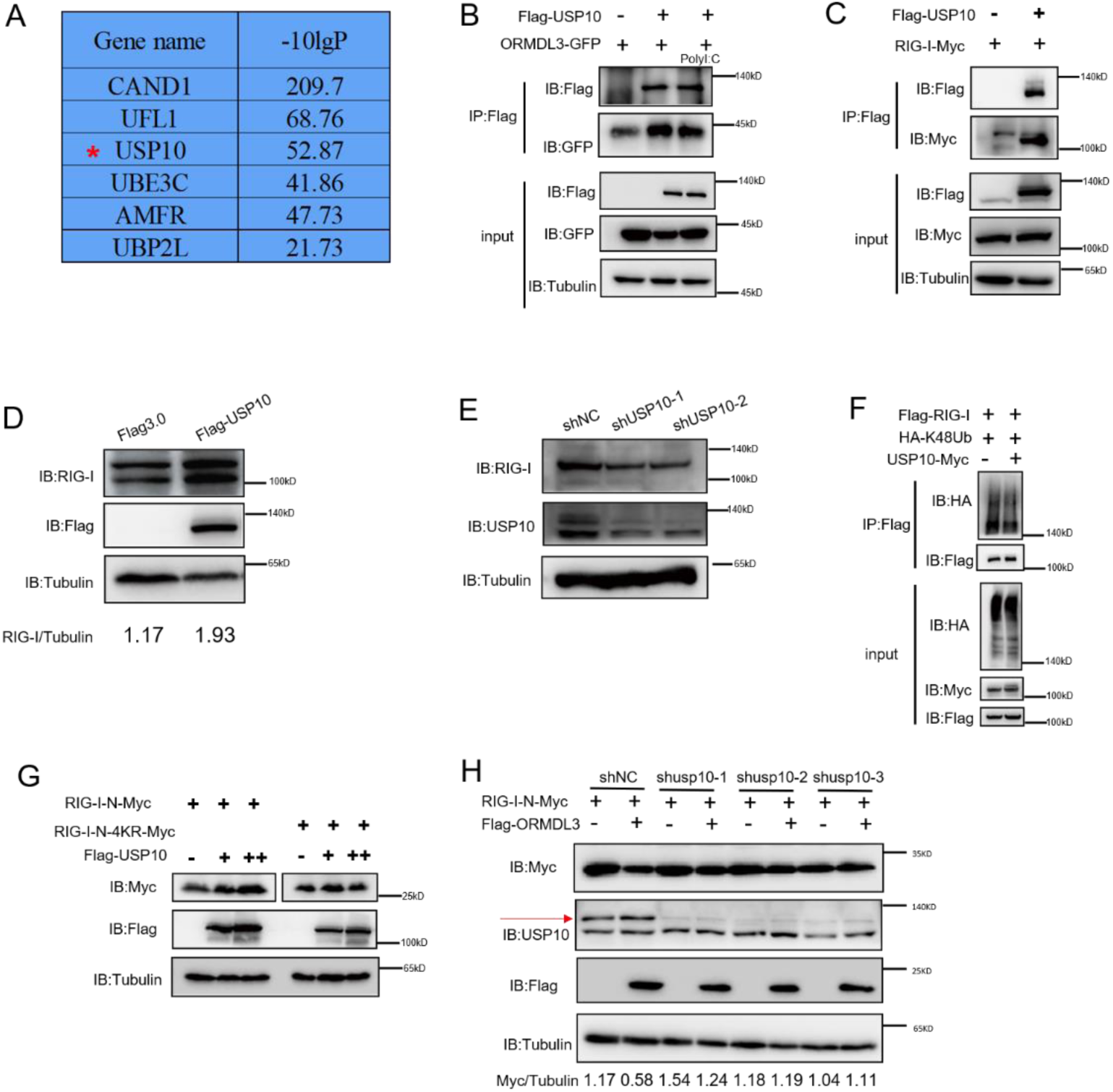
USP10 induces RIG-I stabilization. **(A)** Protein interacted with ORMDL3 screened from mass spectrometry results. **(B)** HEK293T cells were transfected with ORMDL3-GFP and Flag-USP10 plasmids, as indicated, with or without poly(I:C) cotransfection. Cell lysates were immunoprecipitated with the a-Flag antibody, and immunoblotted with a-Flag and a-GFP antibodies. **(C)** HEK293T cells were transfected with RIG-I-MYC and EV or Flag-USP10 plasmids, Cell lysates were immunoprecipitated with the a-Flag antibody, and immunoblotted with a-Flag and a-Myc antibodies. **(D)** Immunoblot the protein level of RIG-I in USP10 stable overexpression HEK293T cell line. **(E)** Immunoblot the protein level of RIG-I in USP10 stable knockdown HEK293T cell line. **(F)** IP and immunoblot analysis of 293T cells transfected with vectors expressing Flag-RIG-I and HA-K48 linked ubiquitin with or without USP10-Myc. **(G)** HEK293T cells were transfected with RIG-I-N-Myc (WT or 4KR) and increasing doses of expression vector for Flag-USP10. The expression levels of RIG-I-N-Myc were analyzed by immunoblot. **(H)** USP10 stable knockdown HEK293T cell line was transfected with RIG-I-N-Myc and Flag-ORMDL3. The expression levels of RIG-I-N-Myc were analyzed by immunoblot.

### ORMDL3 disturbs USP10 induced RIG-I stabilization

Co-immunoprecipitation experiments unveiled robust interactions between USP10 and both RIG-I and ORMDL3, with ORMDL3 demonstrating the ability to disrupt the RIG-I-USP10 interaction **(Figure 6A)**. Crucially, USP10 exhibited a specific role in stabilizing RIG-I, but not other innate proteins such as MAVS, MDA5, and IRF3, and this effect can be reversed by ORMDL3 **(Figure 6B)**. Further investigations showed that the function of ORMDL3 in disturbing USP10-mediated RIG-I stabilization could be rescued by the proteasome inhibitor MG132 but not the lysosome inhibitor CQ **(Figure 6C)**. Subsequent co-transfection experiments delineated that mutations affecting all lysines (KR) or the last nine lysines (mutant1) of RIG-I-N were necessary to prevent USP10-induced accumulation and ORMDL3-mediated degradation **(Figure 6D)**. Notably, single-point mutation of the K146, K154, K164, and K172 residues on RIG-I-N does not affect the regulation of USP10 and ORMDL3 **(Figure 6E)**, while the 4KR mutation abolished this process **(Figure 6F)**. These findings underscored the importance of these four lysine residues in both ORMDL3-mediated degradation and USP10-stabilization of RIG-I. Building upon these observations, we sought to elucidate whether USP10 stabilizes RIG-I through these four sites. Co-expression of HA-K48Ub and RIG-I-N-GFP with or without USP10 revealed a decrease in K48 ubiquitination of RIG-I by USP10, while this effect was nullified in the presence of the 4KR mutant, consistent with ORMDL3-mediated regulation **(Figure 6G)**. Additionally, we verified the functional consequences of USP10-induced RIG-I stabilization by assessing the mRNA levels of *IFNB1, CCL5,* and *ISG54*, which were increased upon enforced USP10 expression and reduced upon co-expression with ORMDL3 **(Figure 6H)**.

**Figure 6.**
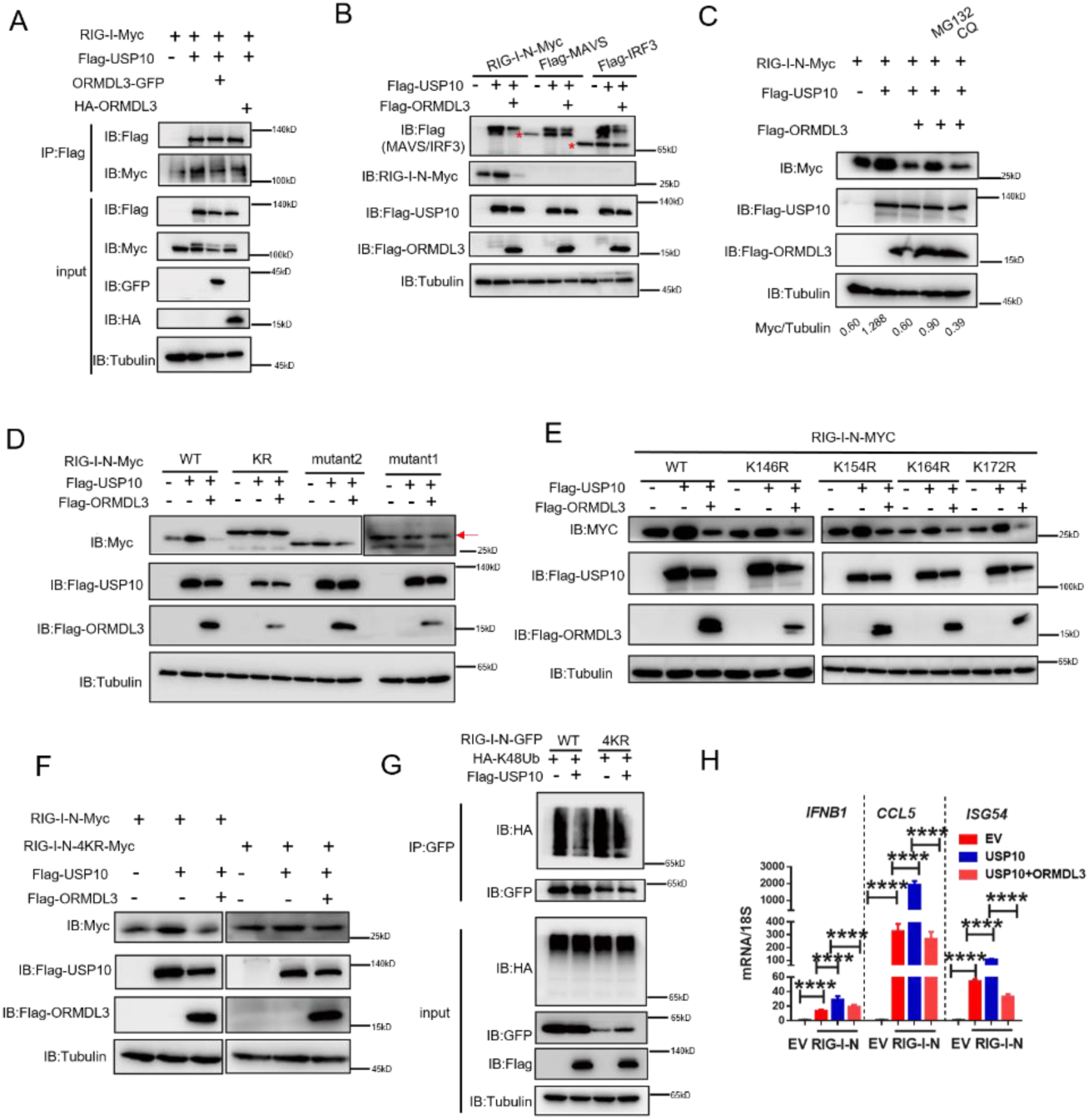
ORMDL3 disturbs USP10-induced RIG-I stabilization. **(A)** HEK293T cells were transfected with Flag-USP10 and RIG-I-Myc plasmids, as indicated, with or without ORMDL3 co-transfection. Cell lysates were immunoprecipitated with the a-Flag antibody, and immunoblotted with a-Flag and a-Myc antibodies. **(B)** Immunoblot analysis of extracts of HEK293T cells transfected with Flag-USP10 and Flag-MAVS and Flag-IRF3 and RIG-I-N-Myc with or without ORMDL3 co-transfection, the cell lysates were analyzed by immunoblot. **(C)** 293T cells were transfected with plasmids encoding RIG-I-N-Myc together with Flag-USP10 with or without Flag-ORMDL3 plasmid treated with MG132 (10 μM), chloroquine (CQ) (50 μM) for 6 h The cell lysates were analyzed by immunoblot. **(D)** HEK293T cells were transfected with Flag-USP10 and RIG-I-N-Myc(WT,KR,mutant1,mutant2) plasmids, as indicated, with or without Flag-ORMDL3 co-transfection. Cell lysates were immunoblotted with a-Flag and a-Myc antibodies. **(E)** 293T cell were transfected with RIG-I-N-Myc (WT or K146R K154R K164R K172R) and Flag-USP10 with or without Flag-ORMDL3. The expression levels of RIG-I-N-Myc(WT or K146R K154R K164R K172R) were analyzed by immunoblot. **(F)** HEK293T cells were transfected with Flag-USP10 and RIG-I-N-Myc(WT and 4KR) plasmids, as indicated, with or without Flag-ORMDL3 co-transfection. Cell lysates were immunoblotted with a-Flag and a-Myc antibodies. **(G)** IP and immunoblot analysis of 293T cells transfected with vectors expressing RIG-I-N-GFP/RIG-I-N-4KR-GFP and HA-K48 Linked ubiquitin with or without USP10 transfection. **(H)** HEK293T cells were transfected with Flag-USP10 and RIG-I-N-Myc and with or without ORMDL3 co-transfection followed by RT-PCR analysis of *IFNB1*,*CCL5*.*ISG54*. *p < 0.05, **p < 0.01, ****p < 0.0001. Data are representative of at least two independent experiments.

### Knock down of ORMDL3 enhance anti-tumor immunity

To assess the impact of ORMDL3 on anti-tumor activity, we conducted knockdown experiments targeting ORMDL3 in LLC and MC38 murine cancer cell lines, followed by subcutaneous inoculation into C57BL/6 mice. Remarkably, the deficiency of ORMDL3 significantly suppressed LCC tumor growth and reduced the tumor formation rate compared to the control group **(Figure 7A-C)**. This tumor growth inhibition by targeting ORMDL3 was further validated in the MC38 cancer model **(Figure 7G-I)**. Moreover, both in LLC and MC38 knocked down cell lines, the protein level of RIG-I was significantly upregulated **(Figure S6A-D)**, and this RIG-I upregulation was further verified by westernblots in LLC tumors **(Figure S6E)** and immunohistochemistry (IHC) in MC38 tumors **(Figure 7K)**. Moreover, investigations revealed that in the LLC tumor model, the knockdown of ORMDL3 led to a significant increase in the expression of ISGs, including *CCL5, CXCL10, TNF*, and *IL-6*, compared to the control group **(Figure 7D)**. This upregulation of ISGs upon ORMDL3 knockdown was consistent in the MC38 cancer model, where *IFNB1*, *CCL5*, and *CXCL10* mRNA levels were significantly elevated **(Figure 7J)**. Flow cytometry analysis demonstrated an increase in CD3^+^ T cell infiltration percentage in LLC tumors with ORMDL3 knockdown **(Figure 7E)**. Notably, although CD8 T cell levels showed no significant change among groups **(Figure S6F)**, activated CD8 T cells (CD8^+^ CD107a^+^ and CD8^+^ CD44^+^) exhibited a remarkable increase in the ORMDL3 knockdown group **(Figure 7F, S6G)**. In addition, IHC assays revealed that more CD8 T cells were infiltrated in ORMDL3 knockdown MC38 tumors **(Figure7K)**. Collectively, these findings suggest that inhibition of tumor-intrinsic ORMDL3 amplifies anti-tumor immunity by increasing ISG expression in the tumor microenvironment (TME) and promoting cytotoxic CD8^+^ T cell activation.

**Figure 7.**
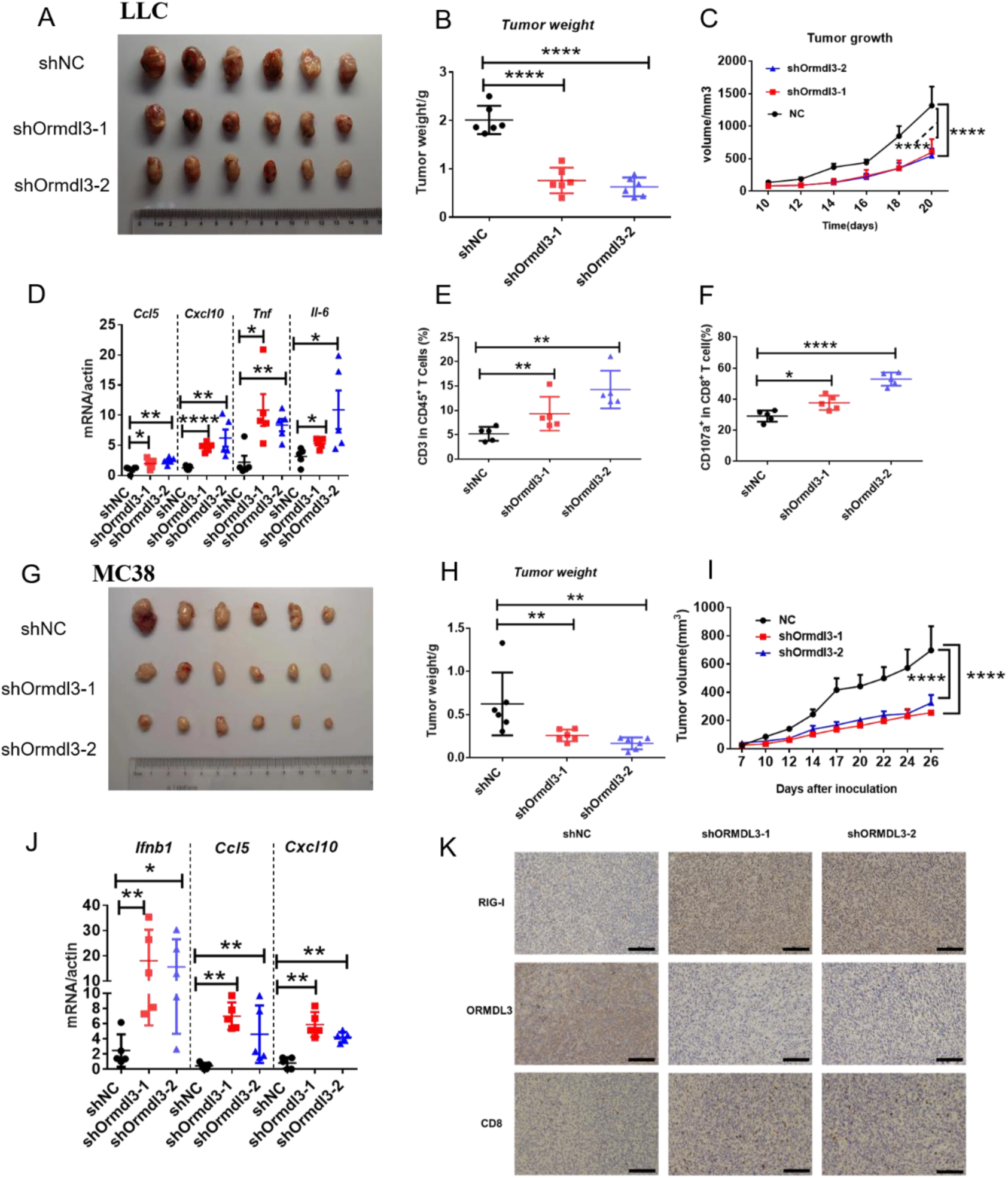
Knockdown of LLC/MC38 ORMDL3 enhances anti-tumor immunity. (A to. **C)** LLC tumor representative images on day 21 after tumor inoculation with shNC, shORMDL3-1, shORMDL3-2 LLC stable knockdown cell line **(A)** tumor weight **(B)** tumor growth **(C)** on day 21 after subcutaneous injection of 1.5×10^^6^LLC in C57BL/6. **(D)** Results of the qRT-PCR assays showing mRNA levels of Ccl5, Cxcl10, Tnf, Il-6 of LLC tumor. **(E,F)** Flow cytometry assay of CD3^+^ T cells in CD45^+^ , CD107a^+^ CD8^+^ T cells percentages. **(G to I)** MC38 tumor representative images on day 27 after tumor inoculation with shNC, shORMDL3-1, shORMDL3-2 MC38 stable knockdown cell line **(G)** tumor weight **(H)** tumor growth **(I)** on day 27 after subcutaneous injection of 5×10^^5^ MC38 in C57BL/6. **(J)** Results of the qRT-PCR assays showing mRNA levels of Ifnb1,Ccl5.Cxcl10 of MC38 tumor. **(K)** Results of the IHC assay showing expression levels of ORMDL3,RIG-I and CD8 in MC38 tumor, scale bars, 50μm.*p < 0.05, **p < 0.01, ****p < 0.0001. Data are representative of at least two independent experiments.

We further analyzed ORMDL3 expression in the TCGA-pan-cancers cohort. We observed higher expression of ORMDL3 in lung adenocarcinoma (LUAD), colon adenocarcinoma (COAD), and lung squamous cell carcinoma (LUSC) compared to their corresponding normal samples **(Figure S7A)**. In LUAD cohorts, high ORMDL3 expression was associated with poor prognosis, as indicated by overall survival (OS), progression-free survival (PFS), and disease-specific survival (DSS) analyses **(Figure S7B-D)**. Analysis of LUAD cohorts also revealed enrichment of stromal scores in tumors with low ORMDL3 expression, Additionally, we found a negative correlation between ORMDL3 expression and the ESTIMATE score, indicating a potential association between ORMDL3 and immune cell infiltration **(Figure S7E)**. Interestingly, ORMDL3 expression showed a negative correlation with CD8 T cell activation markers such as PRF1, GZMA, GZMB, as well as with ISGs such as CCL5 and CXCL10 **(Figure S7F)**, validating our finding that ORMDL3 serves as a negative regulator of the IFN signaling pathway and anti-tumor immunity.

## Discussion

The RIG-I MAVS pathway is essential for the early detection of viral infections and the initiation of an effective antiviral immune response. This pathway has two major downstream signaling events: the intereferon signaling and the pro-inflammatory cytokine signal axis ^43^. During RNA virus infection, MAVS recruits TBK1, which phosphorylates IRF3 and IRF7, leading to the production of type I interferons. Meanwhile, the pro-inflammatory cytokine signaling pathway primarily operates through the activation of NF-κB, resulting in the production of pro-inflammatory cytokines, such as IL-6,TNF, etc ^44^. The activation of RIG-I is critical for innate immunity, and its post-translational modifications play a vital role. For instance, E3 ligases such as TRIM25, TRIM4, RNF135, and RNF194 facilitate RIG-I activation by mediating its K63-linked ubiquitination. Conversely, other E3 ligases, including RNF125, RNF122, casitas B-lineage lymphoma proto-oncogene (c-Cbl, also known as RNF55), and carboxyl terminus of HSC70-interacting protein (CHIP), mediate K48-linked ubiquitination of RIG-I, leading to its degradation through ubiquitin-proteasome pathway and attenuating cascade activation. Additionally, some deubiquitylating enzymes play significant roles in regulating RIG-I activity. For example, USP3, USP21, and CYLD remove K63-linked polyubiquitin chains, thereby repressing RIG-I activity. In contrast, USP4 and USP15 enhance the stability of RIG-I by reducing K48-linked ubiquitylation. Our study discovered that USP10 stabilizes RIG-I by decreasing its K48 ubiquitination at lysine residues146,154,164, and 172.

Research on ORMDL3 has primarily focused on its relationship with asthma and rhinovirus infections. Mechanistic studies have revealed that ORMDL3 regulates intracellular adhesion molecule 1 expression (ICAM1) and modulates ceramide and sphingolipid metabolism ^31^. However, the effects of ORMDL3 on human innate immunity remain to be elucidated further. In our study, we found that the transcription of *IFNB1* and ISGs was significantly impaired in ORMDL3-overexpressing cells in response to viral infection (Figure 1A). Conversely, their transcription induced by RIG-I-N was significantly increased when ORMDL3 was knocked down (Figure 2C). These findings confirm that ORMDL3 is a negative regulator of antiviral innate immunity. ORMDL3 is a multiple transmembrane structural protein located at the core of the serine palmitoyltransferase (SPT) complex, where it stabilizes SPT assembly^41^. Co-IP experiments showed that the endoplasmic reticulum (ER) protein ORMDL3 interacts with mitochondrial protein MAVS, suggesting the existence of scaffold proteins mediating this interaction. ORMDL3 has also been implicated in calcium transport^45,46^. Since calcium transfer between the ER and mitochondria plays an important role in protein synthesis, it is plausible that ER-mitochondria contact site(ERMC) proteins mediate the interaction between ORMDL3 and MAVS. Furthermore, we discovered that the deubiquitinating enzyme USP10 stabilizes RIG-I, while ORMDL3 disturbs this process, thereby negatively regulating IFN production. When USP10 was knocked down, ORMDL3 was unable to degrade RIG-I-N, indicating that USP10 is indispensable for ORMDL3 mediated RIG-I degradation.

Collectively, our findings identify ORMDL3 as a negative regulator of type I interferon pathway and anti-tumor immunity. The proposed working model is that ORMDL3 forms a complex with MAVS and promotes the degradation of RIG-I, thereby attenuating the transcription of type I IFN and cytotoxic CD8^+^ T cell-mediated tumor killing **(Figure S8)**. This negative regulatory loop of antiviral innate immunity mediated by ORMDL3 may provide insights for the development of therapeutics targeting viral infections and tumors.

## Materials and Methods

### Cell lines

HEK293T cell lines (from embryonic kidney of female human fetus), were cultured at 37°C under 5% CO2 in Dulbecco’s modified Eagle’s medium (DMEM) supplemented with 10% fetal bovine serum (ExCell, FSP500), and A549 cell lines (from lung of a 58 years old male human) were cultured in Roswell Park Memorial Institute (RPMI) 1640 medium. LLC and MC38, HCT15, DLD1, SW480, SW620 cell lines were obtained from American Type Culture Collection (ATCC) cultured in Roswell Park Memorial Institute (RPMI) 1640 medium. The cell lines used in this study have been authenticated. Mycoplasma contamination was routinely checked by PCR analysis and eliminated by treatment with PlasmocinTM (ant-mpt). The primers were as follows: Myco forward 5’-GGGAGCAAACAGGATTAGATACCCT-3’; Myco reverse 5’-GCACCATCTGTCACTCTGTTAACCTC-3’.

### Viruses

VSV-GFP was provided by Prof. Rongfu Wang (Zhongshan School of Medicine, Sun Yat-sen University, China) and amplified in Vero cells. HSV-1-GFP was provided by Prof. Musheng Zeng (Sun Yat-sen University Cancer Center, China) and amplified in Vero cells. Cell lines were infected with VSV (0.01 MOI), HSV-1 (0.1MOI) for various times, as indicated in the Figures.

### Plasmid and transfection

Expression plasmids for *RIG-I, MDA5, MAVS, TBK1, TRIF, IKKɛ IRF3* and *IFN-β-luc,ISRE-luc* were provided by Prof. Xin Ye (Microbiology, Chinese Academy of Sciences). Plasmid encoding *ORMDL3* was cloned in pCMV-HA/pCMV-Myc/Pcmv-flag vector, and ORMDL3 truncations ORMDL3(1-42), ORMDL3(43-82), ORMDL3(83-118) and ORMDL3(119-153) were constructed into the pEGFP-N1 vector (CLONTECH Laboratories). USP10 was obtained from prof .Tan (Sun Yat-sen University Cancer Center), and cloned into pCMV-Myc vector. For transfection of plasmids, poly(I:C) (LMW) and poly(dG:dC) (InvivoGen) used in this study into HEK293T, A549 and BMDM cells, DNA Transfection Reagent PEI MW40000, pH 7.1 (Yeasen Biotechnology), Lipo293^TM^ (Beyotime,C0521), Lipofectamine 2000 (ThermoFisher Scientific) or Lipofectamine 3000 (ThermoFisher Scientific) RNAimax (ThermoFisher Scientific) were used according to the manufacturer’s protocol.

### Flow cytometry

Single-cell suspensions were prepared from the tumor tissues of mice, Tumor tissues were cut into small pieces and washed with PBS containing 2% FBS. The tumors were digested in 15 ml RPMI supplemented with 2% FBS, 50 U/ml Collagenase Type IV(Invitrogen, California, USA), 20 U/ml DNase (Roche, Indianapolis, IN) and incubated at 37 °C for 30min to 1 h while gently shaking. Digested tumors were then filtered through a 70 µm strainer after washed three times with PBS. Spleens were mechanically dissociated with a gentle MACS dissociator in RPMI-1640 medium supplemented with 2% FBS. Dissociated spleens were passed through a 70 µm strainer and washed three times with PBS. Red blood cells were lysed for 3 to 5 min with ACK lysis buffer and then washed with PBS containing 2% FBS. Single cells were stained with the appropriate antibodies to surface markers at 4°C for 30 minutes in the dark.The following fluorescent dye-labeled antibodies purchased from BD Biosciences, Biolegend or Invitrogen were used in this study: CD3ε-APC (145-2C11), CD4-Pacific blue (GK1.5), CD8-PE-cy7 (KT15), CD45-APC-cy7 (30-F11), CD44-FITC (IM7), CD107a-PE (1D4B). All flow cytometric data were collected on BD Fortessa X20 (BD Biosciences, San Jose, CA) and performed using Flow-Jo analysis software v10.4. While LLC tumor tissue is grinded into single cell suspension and treat as above described.

### Tumor models

1×10^6^ LLC tumor cells were implanted s.c. into the flanks of mice. 8×10^5^ MC38 tumor cells were implanted the same as LLC tumors, after the tumor was established, measure the volume of tumor once every two days.

### Mass spectrometry and coimmunoprecipitation

1×10^7^ HEK293T cells transfected with flag-vector or flag-ORMDL3 were prepared by washing with cold PBS and then lysed with 1× lysis buffer (Cell Signaling Technology) and incubated on ice for 30 minutes. Supernatants were collected and immunoprecipitated with the indicated antibodies for 4 h at 4 °C, recovered by adding protein A/G Sepharose Beads (Santa Cruz Biotechnology, CA, USA, #sc-2002) overnight. After incubation, beads were washed with wash buffer and immersed in PBS then subjected to mass spectrometry. Immunoblot assays were performed with specific antibodies to identify the proteins interacting with ORMDL3. The following antibodies were used for Co-IP or immunoblot assay: ORMDL3 (abcam,107639) (abcam 211522), Flag (Sigma, St. Louis, USA, #F1804), HA (Beijing Ray Antibody Biotech, RM1004). The rabbit antibody GAPDH (1:5000 for immunoblot, #GB15004) and rabbit antibody beta-Actin (1:2000 for immunoblot, #GB15003) were purchased from Servicebio Biotechnology (Wuhan, China). Rabbit anti-GFP (50430-2-AP), mouse anti-GFP (66002-1-Ig), anti-β-tubulin (66009-1-Ig), rabbit anti-myc(10828-1-AP), mouse anti-myc(60003-2-Ig) were ordered from Proteintech. Mouse anti-RIG-I (Santa Cruz, sc376845), rabbit anti-USP10 (ABclona, A7505). Secondary antibodies were purchased from The Jackson Laboratory. The cell lysis buffer (10 × ) (#8903) was bought from Cell Signaling Technology. Protein Marker (DB180-10) was bought from MIKX. The protease inhibitor cocktail and phosphatase inhibitor cocktail were purchased from TargetMol (C0001).

### Dual-luciferase reporter assay

Cells were transfected with plasmids encoding IFN-β or ISRE luciferase reporter and RIG-I(N), MDA5, MAVS, TBK1, IRF3-5D together with pRL-TK and the plasmids encoding ORMDL3. Cells were collected and lysed 24 h post-transfection. Subsequently, the luciferase activities were measured using a Dual-luciferase Reporter Assay System (Promega, Madison, USA, #E1910). Normalization of data by the ratio of firefly luciferase activity to renal luciferase activity. Each group was measured in triplicate.

### RNAi

All the siRNA oligonucleotides containing 3′dTdToverhanging sequences were chemically synthesized in genepharma (Suzhou, China) and transfected into cells using Lipofectamine ™ RNAiMAX Transfection Reagent (Thermo Fisher).The siRNAs corresponding to the target sequences were synthesized in RIBOBIO (Guangzhou, China). In this study, the siRNAs sequences were designed as follows: ORMDL3 si #1, 5′ - GCAUCUGGCUCUCCUACGUTTdTdT 3′; ORMDL3si #2, 5′ - GGCAAGGCGAGGCUGCUAATTdTdT 3′; ORMDL3 si #3, 5′-

CCCUGAUGAGCGUGCUUAUTTdTdT 3′. For transfection of siRNAs used in this study into HEK293T cells, Transfection Reagent Lipofectamine RNAiMAX (ThermoFisher Scientific) was used according to the manufacturer’s protocol.

### RNA extraction and quantitative real-time PCR

Total RNA from cells was extracted with Trizol reagent (Vazyme R711) according to the manufacturer’s instructions. Tumor tissue RNA was grinded by Sample freezing grinder: LUCA(LUKYM-I). Complementary DNA was synthesized using the HiScript II Q RT SuperMix (Vazyme, R223-01). SYBR Green Mix (GenStar, A301-10) was used for quantitative real-time PCR (qRT-PCR) assays.. Relative quantification was performed with the 2(-ΔΔCT) method using 18S (for human cells) or Actb (β-Actin, for mouse cells) for normalization. The Specific qRT-PCR primers are listed in Supplementary Table 1.

### Establishment of overexpressed stable cells and knock down cell lines

ORMDL3 cDNA was constructed into the pCDH-CMV-MCS-EF1 vector. A549 and HEK293T cells which stably overexpress plasmid encoding ORMDL3 were generated by lentivirus-mediated gene transfer. HEK293T cells were co-transfected with lentiviral expressing plasmid, lentiviral packaging plasmid psPAX2 (Addgene, Cat#12260) and VSV-G envelope expressing plasmid pMD2.g (Addgene, Cat#12259). After 48 hours, the lentiviruses were used for infecting A549 cells and then selected cells with puromycin (ThermoFisher Scientific). Human ORMDL3 knockdown cell line:shORMDL3 -1 and shORMDL3-2.Human USP10 knockdown cell line: shUSP10-1 and shUSP10-2, shUSP10-3.Mice Ormdl3 knockdown cell line shOrmdl3-1 and shOrmdl3-2.annealing oligos were ligated into PLKO.1 vector and after virus package infected target cell and then selected with puromycin. the shRNA sequences are listed as follows. shORMDL3-1: CCGGCCCACAGAATGTGATAGTAATCTCGAGATTACTATCACATTCTGTGGGTTTTTG ; shORMDL3-2 : CCGGCATGGGCATGTATATCTTCCTCTCGAGAGGAAGATATACATGCCCATGTTTTTG ; shUSP10-1: CCGGCCTATGTGGAAACTAAGTATTCTCGAGAATACTTAGTTTCCACATAGGTTTTTG ;shUSP10-2: CCGGCCCATGATAGACAGCTTTGTTCTCGAGAACAAAGCTGTCTATCATGGGTTTTTG ; shUSP10-3: CCGGCGACAAGCTCTTGGAGATAAACTCGAGTTTATCTCCAAGAGCTTGTCGTTTTTG ;shOrmdl3-1: CCGGCCAAGTATGACCAAGTCCATTCTCGAGAATGGACTTGGTCATACTTGGTTTTTG ;shOrmdl3-2: CCGGGCCGACTTGGAGTAGCTTGTACTCGAGTACAAGCTACTCCAAGTCGGCTTTTTG.

### BMDMs

Macrophages were differentiated from the bone marrow of wild-type (WT) C57BL/6 mice All bone marrow cells were flushed out and filtered through a 70-µm cell strainer. After centrifugation, red blood cells were lysed. The resultant bone marrow cells were resuspended in RPMI 1640 (Gibco) supplemented with 10% FBS (Gibco), 1% penicillin/streptomycin (Gibco), and 50 μM 2-mercaptoethanol (Sigma) in the presence of 20 ng /ml macrophage colony-stimulating factor (M-CSF; PeproTech) for 7 days and mature BMDMs were stimulated with indicated stimulation poly(I:C) or poly(dG:dC).

### AAV virus production

All AAV vectors were produced in HEK293 cells via the triple plasmid transient transfection methods as pAdltea:Paav2/1:target gene=750ng:450ng:375ng. For small-scale preps, HEK293T cells were seeded in 10-cm dishes and grown to 80% confluence in Dulbecco’s modified Eagle’s medium (DMEM) containing 10% fetal bovine serum (FBS) (Gibco, 26140079) and 1% PenStrep (Thermo FisherScientific, 15140122). Cells were then triple transfected with the vector pscAAV-CAG-GFP (Addgene, 83279) or pscAAV-CAG-Ormdl3, AAV1 Rep/Cap (Addgene, 112862), and Ad helper plasmid (pAddelta F6 from Addgene, 112867) at a ratio of 1:1.2:2 (3.75:4.5:7.5 μg per 10-cm dish) using PEI MW40000, pH 7.1 (Yeasen Biotechnology, 40816ES03 at a ratio 4:1 of PEI/total DNA. Cells were harvested 3 days post transfection by scraping cells off the plate in their conditioned medium and lysing cells through 3× freeze-thaw cycles between 37°C and - 180°C. Preps from three replicate plates were then pooled, incubated with 25 U/mL of benzonase (Millipore Sigma, 20 E8263-25KU) at 37°C for 1 h to remove plasmid and cell DNA, centrifuged at 4°C and 1 4,000 × g for 30 min, and the supernatant filtered through a 0.22-μm polyethersulfone (PES) bottle-top filter (Corning, 431097). The filtered lysate was Purificated by iodixanol gradient ultracentrifugation. For AAV collection, the fractions obtained from the 40% phase were analyzed by measuring absorbance at 20-fold dilution at 340 nm to identify the main contaminating protein peak, as previously described. For ultrafiltration/concentrated AAV, 0.001% Pluronic F68 +200mM NaCl PBS was added to the pool to reach a total volume of 15 ml, using Amicon Ultra-15 centrifugal filter units (MWCO, 100 kDa; Merck Millipore). After concentration to a minimum of 500 µl, the product was aliquoted and stored at -80°C.

### AAV titration

Prepare a plasmid stock of 2×10 molecules/μl to generate a standard curve, and then treat the puri¦ed AAV samples with DNase I to eliminate any contaminating plasmid DNA carried over from the production process (DNase does not penetrate the virion). Make 6 serial dilutions of reference sample and DNase-treated and AAV samples and detect with RT-QPCR. And then perform data analysis using the instrument’s software. Determine the physical titer of samples (viral genomes (vg)/mL) based on the standard curve and the sample dilutions.

### Immunofluorescence labeling and confocal microscopy For FRET assay

YFP-MAVS and CFP-ORMDL3 expressing plasmids were co-transfected into HeLa cells and incubated for 24h. Cells were fixed with 4% paraformaldehyde for 30 min, Images were observed on laser confocal fluorescence microscopy (Zeiss, LSM880), we bleached YFP-MAVS and measured the MFI of YFP-MAVS and CFP-ORMDL3.

### Co-immunoprecipitation and immunoblot analysis

For immunoprecipitation (IP), cells were lysed with lysis buffer (Cell Signaling Technology) supplemented with protease inhibitor for 30 min. After centrifugation at 12,000 rpm and 4°C for 10 min, supernatants were collected and incubated with appropriate antibodies for 1 h and protein G beads (Santa Cruz Biotechnology) overnight. Thereafter, the beads were washed four times with cold PBS, followed by SDS-PAGE and immunoblot analysis. For immunoblot analysis, cells or tissues were lysed with RIPA buffer (Cell Signaling Technology). Protein concentrations were measured with Bradford Protein Assay Kit (Beyotime), and equal amounts of lysates were used for SDS-PAGE. The samples were eluted with SDS loading buffer by boiling for 10 min and then performed SAS-PAGE. The proteins were transferred onto PVDF membrane (Roche), and immunoblot analysis was performed with appropriated primary antibodies at 4°C overnight (Geng et al, 2017) and horseradish peroxidase (HRP) conjugated secondary anti-mouse or anti-rabbit antibodies for 1 h at room temperature. ChemiDoc Touch (Bio-Rad) achieved visualization.

### Statistical analysis

The data were analyzed with GraphPad Prism 7. For two independent groups, the student’s t test was used to determine statistical significance. Statistical details for individual experiments can be found in the Figure legends. Statistical significance was two-tailed and p < 0.05 is considered statistically significant *P value*s are indicated by asterisks in the Figures as follows: *p < 0.05, **p < 0.01, ***p < 0.001, and n.s. indicates non-significant.

### Ethics Approval and Consent to Participate

All animal studies in this study were carried out according to the National Institute of Health Guide for the Care and Use of Laboratory Animals with the approval of Sun Yat-Sen University Cancer Center Institutional Animal Care and Use Committee (approval number: SYSU-IACUC-2023-000104).

## Acknowledgements

This project was supported by grants from the National Natural Science Foundation of China (82273045) and the Guangdong Basic and Applied Basic Research Foundation (2022A1515011930).

## Author Contribution

Q.Z performed most of the experiments and analyses. C.Y and Q.Z performed the mouse experiments. S.C conceived the study. C.S, S.Z, Y.M, J.W and Z.W provided technical assistance. S.C and C.S supervised the study. S.C and Q.Z wrote the manuscript.

## Data Availability Statement

Raw data that support the findings of this study has been deposited in Research Data Deposit database (http://www.researchdata.org.cn) with the Approval Number RDDB2024887515. Any reasonable requests for this study are available from the corresponding author.

## Supplementary materials

This supplementary materials includes

1) Seven supplementary Figures
2) One supplementary Table

**Supplementary Figure 1.**
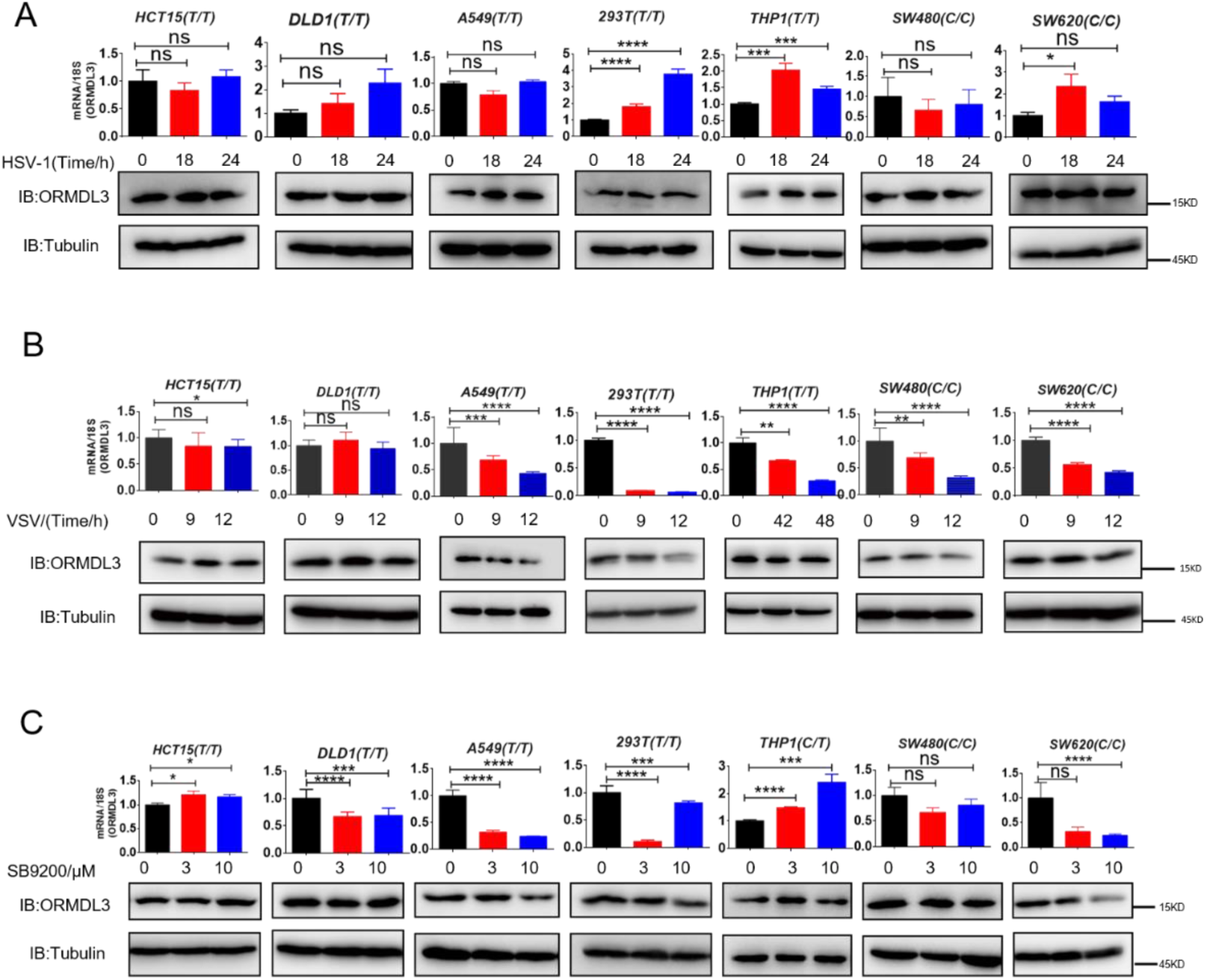
ORMDL3 expression under different treatments. **(A)** Results of the WB and qRT-PCR assays showing ORMDL3 protein and RNA levels HCT15 DLD1 SW480 SW620 CRC cell line and A549 293T THP1 infected with HSV (MOI = 0.1) for indicated times. **(B)** Results of the WB and qRT-PCR assays showing ORMDL3 protein and RNA levels HCT15 DLD1 SW480 SW620 CRC cell line and A549 293T THP1 infected with VSV-1 (MOI = 0.01) for indicated times. **(C)** Results of the WB and qRT-PCR assays showing ORMDL3 protein and RNA levels HCT15 DLD1 SW480 SW620 CRC cell line and A549 293T THP1 treated with RIG-I agonist SB9200 for indicated times.

**Supplementary Figure 2.**
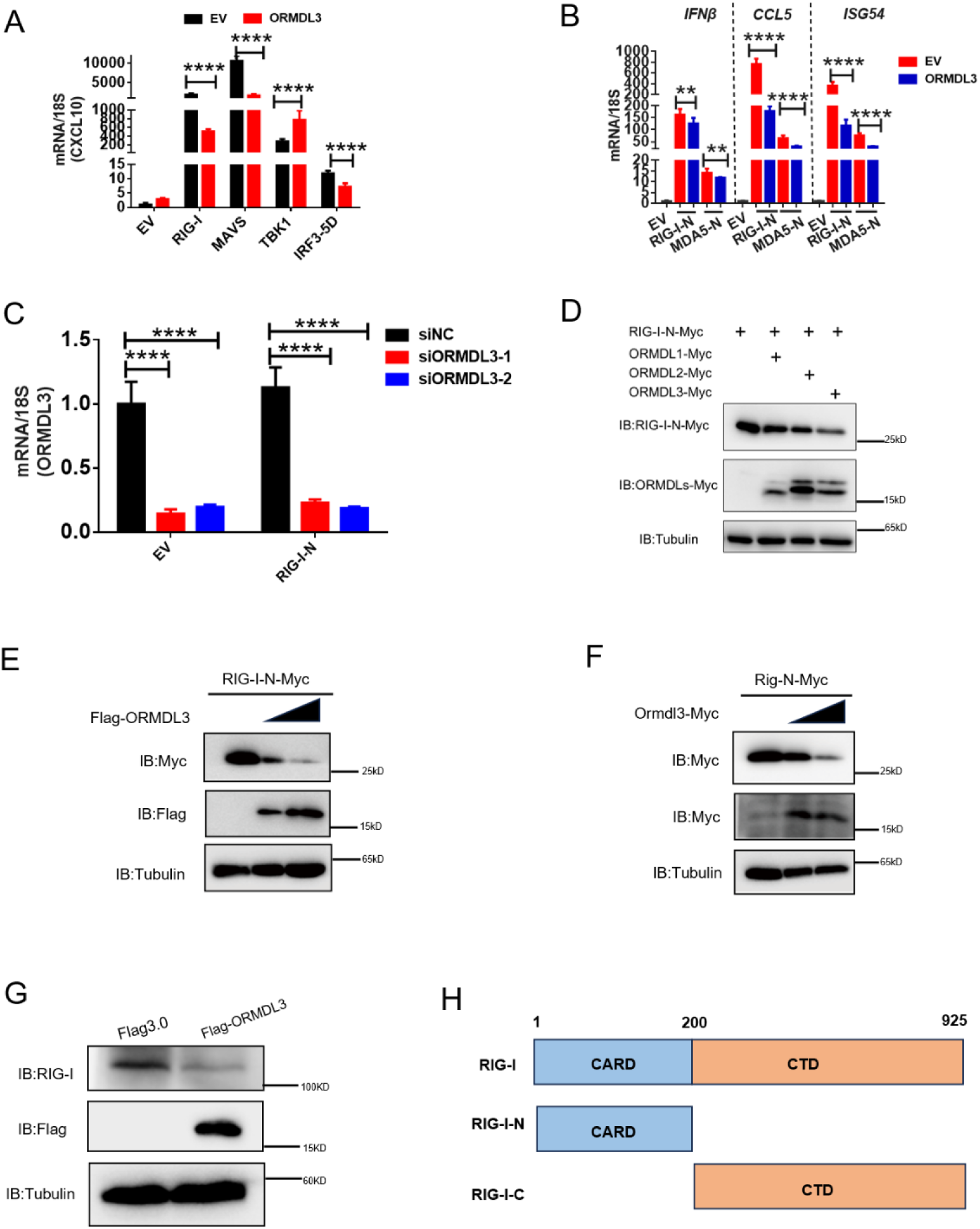
ORMDL3 downregulation is conservative in both human and murine cells. **(A)** RT-PCR analysis of CXCL10 when ORMDL3 was co-expressed with RIG-I MAVS TBK1 or IRF3-5D. **(B)** Results of the qRT-PCR.assays showing IFNB1, CCL5, ISG54 mRNA in HEK293T cells transfected with EV or ORMDL3 plasmids together with individual RIG-I-N or MDA5-N plasmids for 24 h. **(C)** Results of the qRT-PCR.assays showing ORMDL3 knockdown efficiency which links to figure 2D. **(D)** Immunoblot assay of RIG-I-N expression in HEK293T co-transfected with ORMDLs families. **(E)** Immunoblot assay of exogenous RIG-I expression in HEK293T transfected with human ORMDL3. **(F)** Immunoblot assay of exogenous Rig-I expression in HEK293T transfected with mice Ormdl3. **(G)** Immunoblot assay of endogenous RIG-I expression in HEK293T transfected with Flag-ORMDL3. **(H)** The diagram of RIG-I truncations.

**Supplementary Figure 3.**
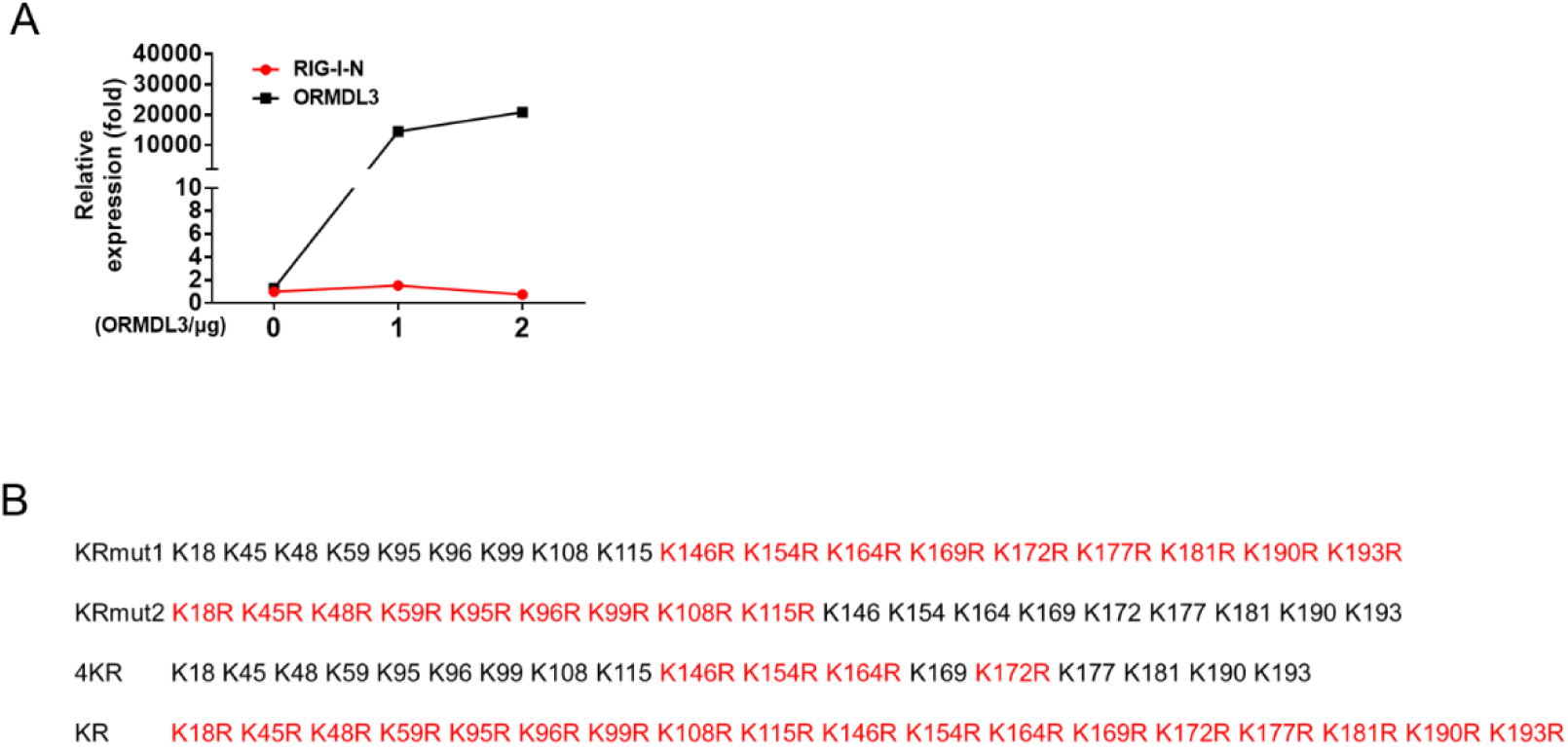
ORMDL3 promotes the degradation of RIG-I. **(A)** RT-PCR assay of RIG-I-N mRNA levels in HEK293T cells transfected with RIG-I-N and increasing amounts of ORMDL3. **(B)** Different annotations of KR mutation of RIG-I-N.

**Supplementary Figure 4.**
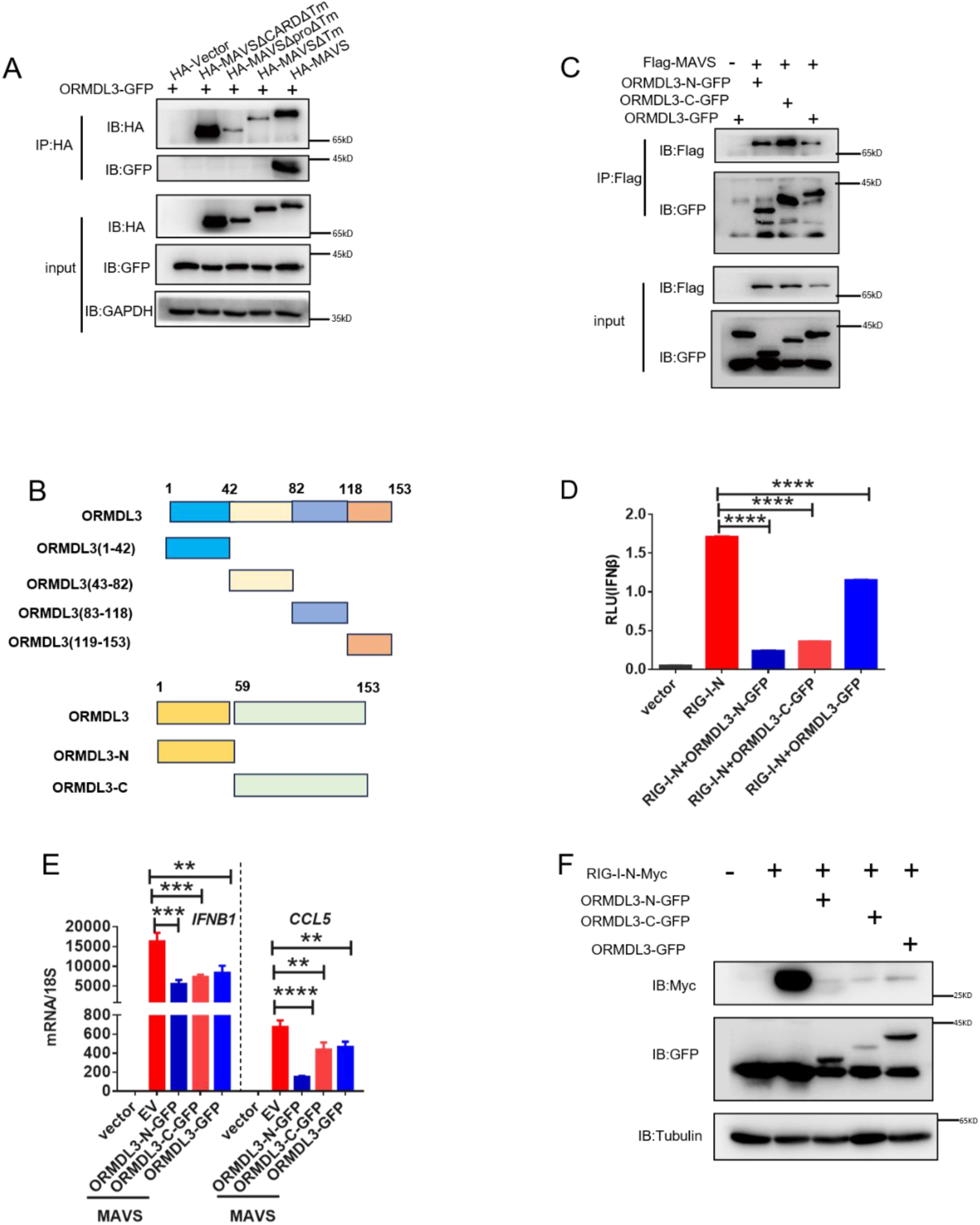
All truncations of ORMDL3 can inhibit the interferon production. **(A)** HEK293T cells were transfected with the indicated plasmids, and cell lysates were immunoprecipitated with an HA antibody (a-HA) followed by immunoblots using GFP (a-GFP) and a-HA antibodies. **(B)** Scheme of the truncations of ORMDL3. **(C)** HEK293T cells were transfected with the indicated plasmids, and cell lysates were immunoprecipitated with a Flag antibody (a-Flag) followed by immunoblots using GFP (a-GFP) and a-Flag antibodies. **(D)** Results of the luciferase assays showing IFNβ activity in HEK293T cells transfected with ORMDL3 truncations with RIG-I-N. **(E)** Results of the qRT-PCR assays showing mRNA levels of IFNB1, CCL5 in HEK293T cells transfected with ORMDL3 truncations with MAVS. **(F)** HEK293T cells were transfected with RIG-I-N-Myc and different truncation of ORMDL3 and followed by immunoblot assay.

**Supplementary Figure 5.**
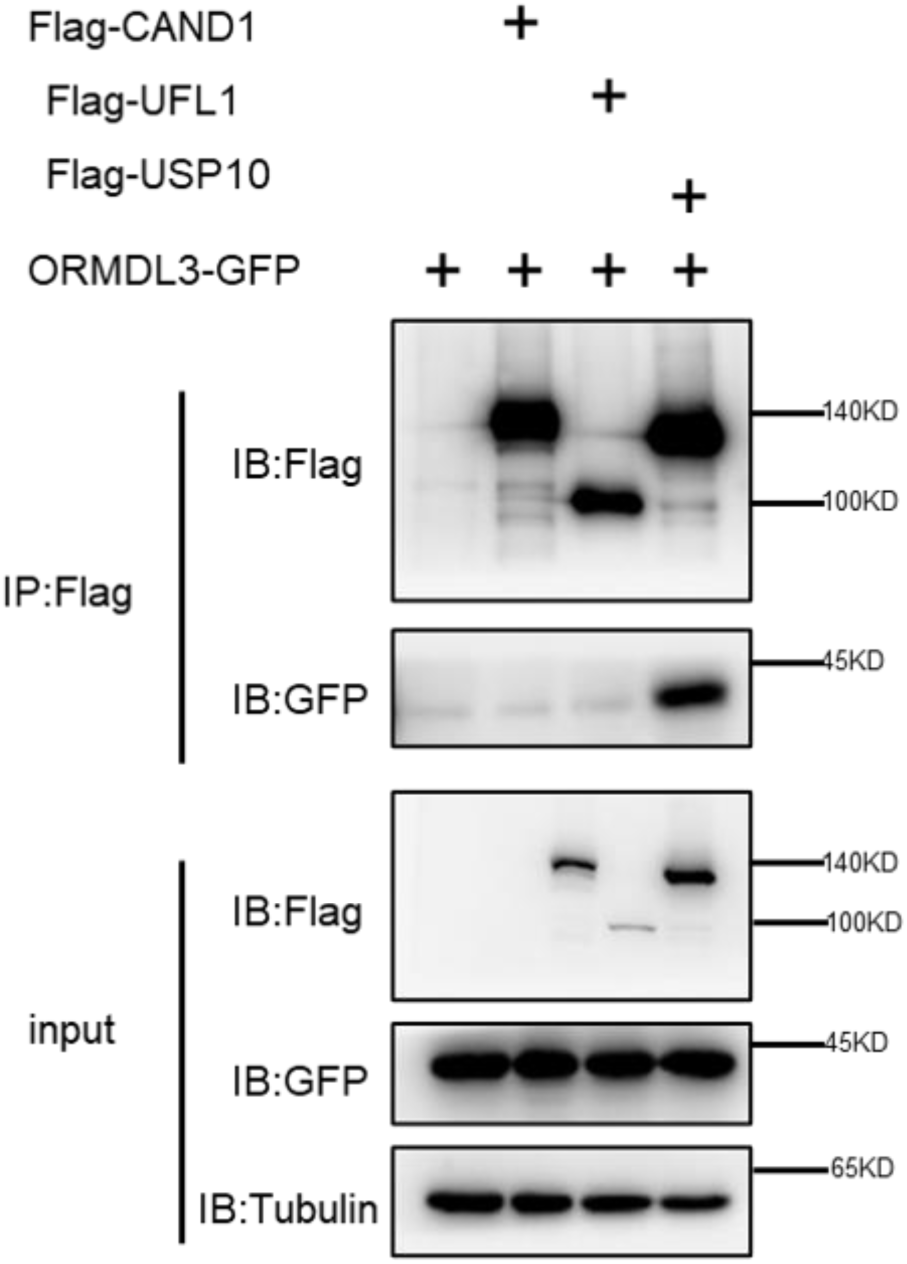
ORMDL3 interacts with USP10 but not CAND1 or UFL1. HEK293T cells were transfected with the indicated plasmids, and cell lysates were immunoprecipitated with an Flag antibody (a-Flag ) followed by immunoblots using GFP (a-GFP) and Flag (a-Flag) antibodies.

**Supplementary Figure 6.**
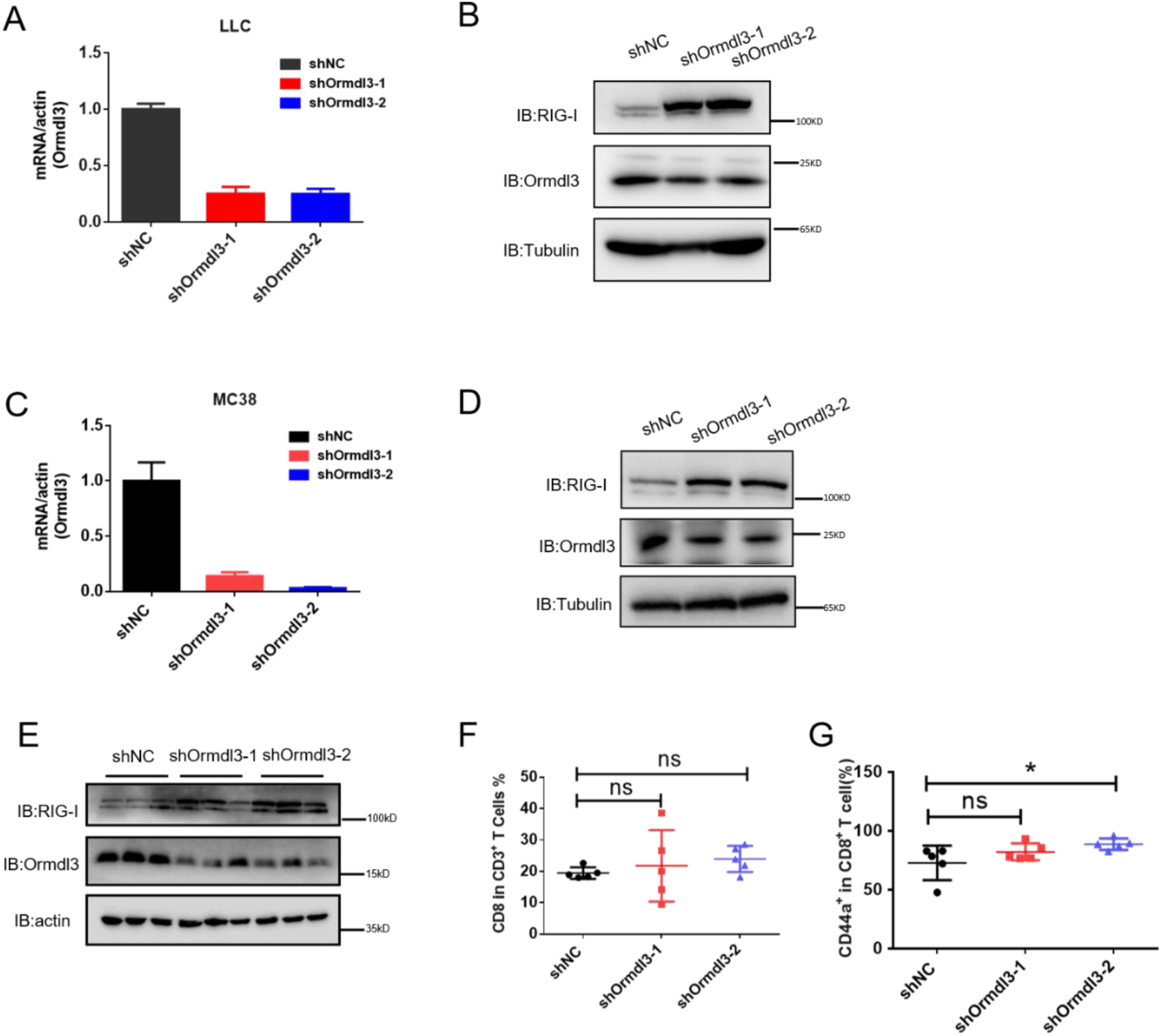
Inhibiting ORMDL3 enhances the abundance of RIG-I. **(A)** RT-PCR showing the shRNA knockdown of ORMDL3 or shRNA control in LLC lung cancer cells. **(B)** Immunoblot showing the shRNA knockdown of ORMDL3 or shRNA control in LLC lung cancer cells. **(C)** RT-PCR showing the shRNA knockdown of ORMDL3 or shRNA control in MC38 colon cancer cells. **(D)** Immunoblot showing the shRNA knockdown of ORMDL3 or shRNA control in MC38 colon cancer cancer cells. **(E)** Immunoblot assay of LLC tumor in group shNC, shOrmdl3-1,shOrmdl3-2, followed by analysis of protein level of ORMDL3 and RIG-I. **(F,G)** Flow cytometry assay of CD8^+^T cells in CD3^+^ and CD44^+^CD8^+^ T cells percentages.

**Supplementary Figure 7.**
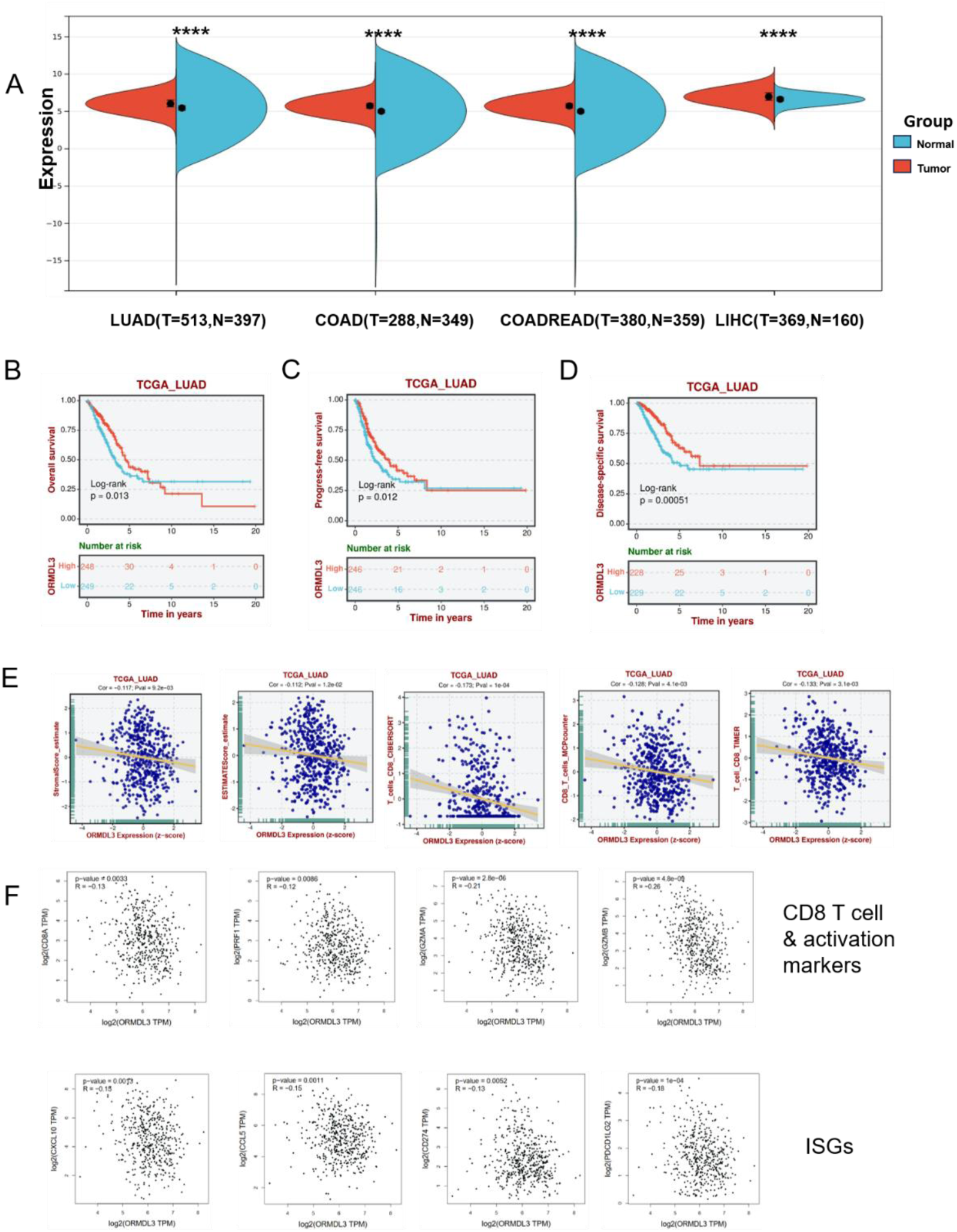
ORMDL3 expression is associated with poor survival and reduced immune cell infiltration and ISGs. **(A)** ORMDL3 expression between tumor and adjacent normal tissues in The Cancer Genome Atlas(TCGA) pan-cancer cohorts. **(B to D)** Association of ORMDL3 level with overall survival(OS),progression free survival(PFS) and disease-specific survival(DSS) in TCGA-LUAD. **(E)** Correlation among ORMDL3 expression with stromalscore and CD8T cell infiltration in LUAD cohort from TCGA datasets. **(F)** Correlation among ORMDL3 expression with CD8 activation markers: CD8A, PRF1, GZMA, GZMB and ISG, CCL5, CXCL10, CD274 and PDCD1LG2 in The Cancer Genome Atlas (TCGA) pan-cancer cohorts from the TIMER2.0 database.

**Supplementary Figure 8.**
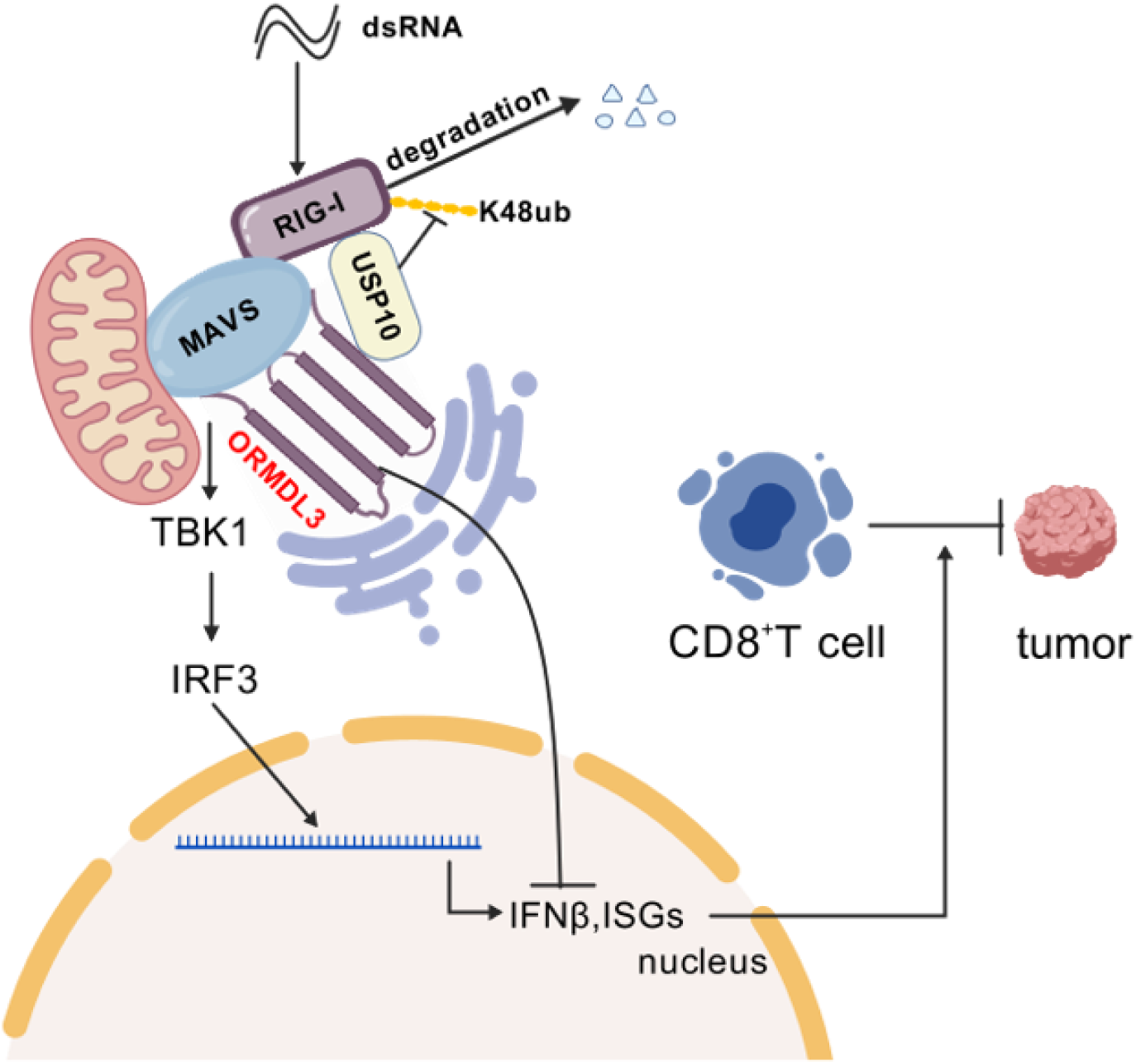
**Schematic shows that ORMDL3 promotes the degradation of RIG-I and attenuates IFN in cancer.**

**Supplementary Table 1.**
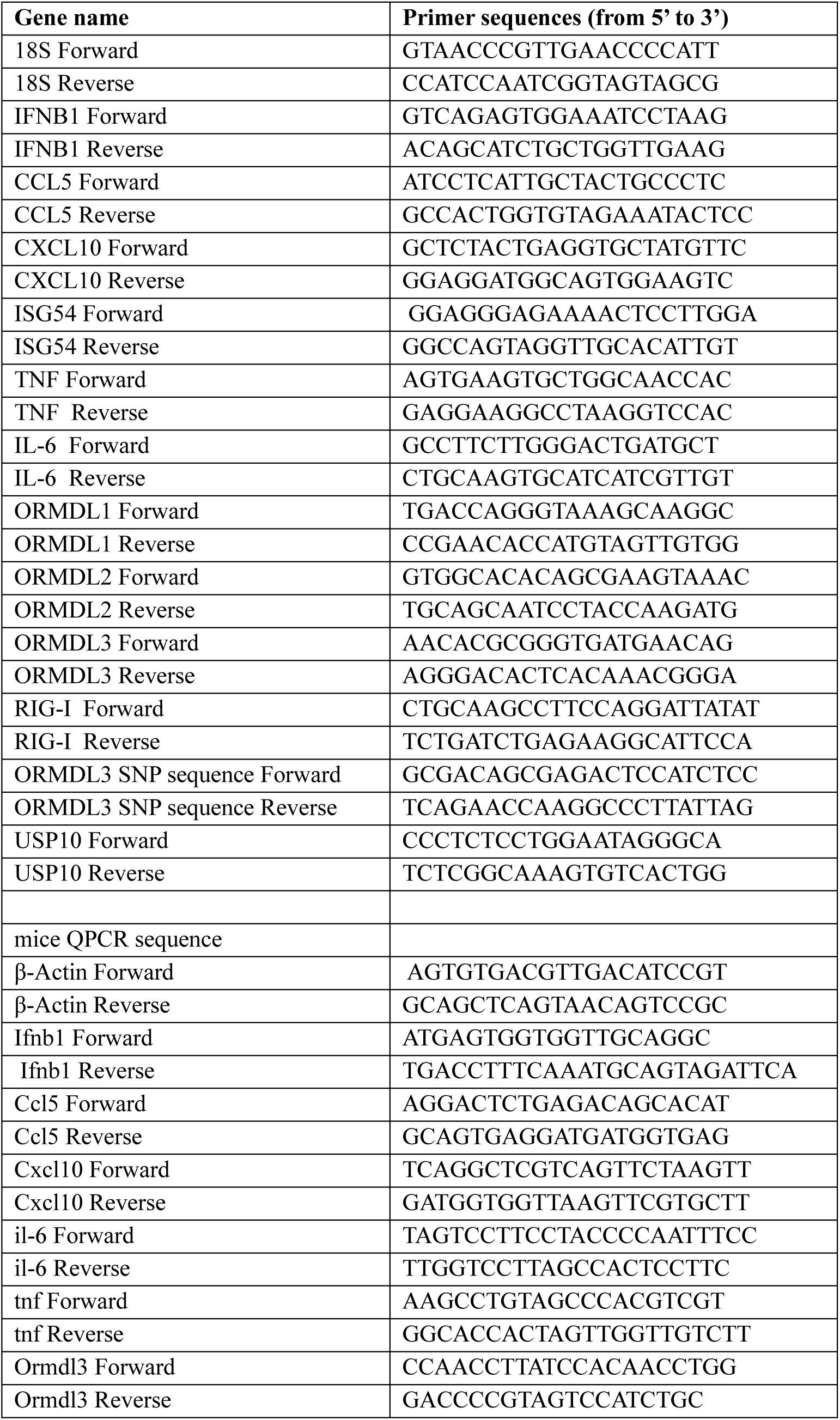
Primers for qPCR. Related to Materials and Methods.

